# Interplay between chemotaxis, quorum sensing, and metabolism regulates *Escherichia coli*-*Salmonella* Typhimurium interactions *in vivo*

**DOI:** 10.1101/2024.08.19.608533

**Authors:** Leanid Laganenka, Christopher Schubert, Manja Barthel, Bidong D. Nguyen, Jana Näf, Thomas Lobriglio, Wolf-Dietrich Hardt

**Affiliations:** Institute of Microbiology, D-BIOL, ETH Zurich, Switzerland

**Author notes:** These authors contributed equally to this work (Leanid Laganenka, Christopher Schubert).

## Abstract

Motile bacteria use chemotaxis to navigate complex environments like the mammalian gut. These bacteria sense a range of chemoeffector molecules, which can either be of nutritional value or provide a cue for the niche best suited for their survival and growth. One such cue molecule is the intra- and interspecies quorum sensing signaling molecule, autoinducer-2 (AI-2). Apart from controlling collective behavior of *Escherichia coli*, chemotaxis towards AI-2 contributes to its ability to colonize the murine gut. However, the impact of AI-2-dependent niche occupation by *E. coli* on interspecies interactions *in vivo* is not fully understood.

Here, using the C57BL/6J mouse infection model, we show that chemotaxis towards AI-2 contributes to nutrient competition and thereby affects colonization resistance conferred by *E. coli* against the enteric pathogen *Salmonella enterica* serovar Typhimurium (*S.* Tm). Like *E. coli*, *S.* Tm also relies on chemotaxis, albeit not towards AI-2, to compete against residing *E. coli* in a gut inflammation-dependent manner. Finally, by using a barcoded mutant library pool of *S.* Tm, we analyzed how AI-2 signaling in *E. coli* affects the central metabolism of *S.* Tm. AI-2-dependent niche colonization by *E. coli* specifically affected the fitness of *S.* Tm mutants deficient in fumarate respiration (Δ*dcuABC*) or mannose (Δ*manA*) utilization. Our findings thus provide important insights into AI-2-mediated *E. coli*-*S.* Tm interactions during gut infection.

**Author Summary:** Both chemotaxis and AI-2 quorum sensing systems have been extensively studied in *Escherichia coli*. Despite our understanding of these systems at a molecular level *in vitro*, their physiological relevance *in vivo*, particularly in the context of mammalian gut colonization, remains less explored. Building on our previous work on the role of chemotaxis and AI-2 signaling in *E. coli* gut colonization, we investigated their roles in interspecies interactions. Specifically, we examined how AI-2-dependent colonization by *E. coli* affects its competition with the enteric pathogen *Salmonella enterica* serovar Typhimurium (*S.* Tm) and the metabolic requirements for *S.* Tm growth.

Our data show that AI-2 signaling contributes to colonization resistance of *E. coli* against *S.* Tm. Although *S.* Tm also requires chemotaxis to grow efficiently in *E. coli*-colonized mice, this is independent of its ability to sense AI-2. Notably, AI-2-dependent niche occupation by *E. coli* altered *S.* Tm metabolism at different stages of infection. Collectively, our findings highlight how AI-2 signaling in one species can affect the metabolism of its interaction partners *in vivo*.

## Introduction

Chemotaxis systems allow motile bacteria to navigate environmental gradients of various chemical compounds to locate niches that are preferable for their survival and growth [1–3]. Although originally studied in context of single cell behavior, the role of chemotaxis in mediating bacteria-bacteria and bacteria-host interactions is now becoming increasingly evident [4]. Importantly, although encoded by a minority of host-associated bacteria, including a few strains of the normal gut microbiota, motility and chemotaxis systems are more common among bacterial pathogens [5]. In these bacteria, chemotaxis has been shown to be an important factor in host colonization and development of disease. The examples include, but are not limited to human pathogens infecting diverse body sites: *Helicobacter pylori*, *Pseudomonas aeruginosa*, *Vibrio cholerae*, *Borrelia burgdorferi* and *Salmonella enterica* serovar Typhimurium [6–12]. The latter, a causative agent of acute gastroenteritis, requires chemotaxis for gut colonization and colitis development in the mouse model [13,14]. However, the role of chemotaxis in host colonization is not solely associated with pathogenic bacteria. Host colonization by commensal *Vibrio fischeri*, *E. coli* and *Lactobacillus agilis* strains was shown to be enhanced by motility and chemotaxis as well [15–18].

Although bacteria mainly rely on chemotaxis to detect and reach the sources of compounds with certain nutritional value, it is not always the case during host colonization. In this context, host- or resident microbiota-produced cues serve as a guiding signal to direct colonizing bacteria towards their respective niches [4,19]. Urea chemotaxis and pH sensing has been implied in *H. pylori* infection [20,21], whereas chemotaxis towards host-produced mucus and hormone norepinephrine promotes gut colonization by *V. cholerae* and pathogenic *E. coli*, respectively [22,23]. In our previous study, we have further identified interspecies quorum sensing (QS) molecule autoinducer-2 (AI-2) as a chemotactic signal that promotes gut colonization by commensal *E. coli* strains [18]. AI-2 is produced and sensed by a variety of bacteria, with AI-2 mimics being produced by epithelial cells and *Saccharomyces cerevisiae in vitro* [24,25]. This allows AI-2 to control the crosstalk and coordination of collective behaviour on interspecies and potentially interdomain levels. Importantly, AI-2 is a major autoinducer molecule in the mammalian gut. Manipulation of luminal AI-2 concentration influences the abundance of the major bacterial phyla of the gut microbiota, which in turn might affect its function [26,27]. However, the molecular nature of such interspecies or interphylum effects is not fully understood.

Although chemotaxis allows host-colonizing bacteria to effectively locate the most suitable niche, they also require certain metabolic capabilities for efficient growth within that niche. In the case of enteropathogen colonization, there is also a transition from a healthy gut to an inflamed state, which causes a significant alteration in the gut luminal environment. This change introduces various inorganic electron acceptors, such as oxygen, nitrate, and tetrathionate, which promote the proliferation of facultative anaerobic bacteria [28–30]. Seminal studies on *E. coli* have elucidated their carbohydrate preferences for both commensal and pathogenic strains to colonize the murine model [31,32]. Additionally, D-galactitol has been identified as a key factor in both intra- and interspecies competition between *S.* Tm and *E. coli* [33,34]. Furthermore, AI-2, albeit a quorum sensing molecule, is tightly integrated into the cellular metabolism. It is produced as a byproduct of the activated methyl cycle, and AI-2-mediated quorum sensing may be integrated into carbon catabolite repression (CCR) through an interaction between the AI-2 kinase LsrK and HPr, a component of the phosphotransferase system [24,35]. The expression of the *lsr* operon, which is required for AI-2 import (*lsrACDB*), degradation (*lsrFG*) and chemotaxis (*lsrB*), is likewise subject to CCR via cAMP receptor protein (CRP) [36,37]. The link between quorum sensing and CCR offers a compelling mechanism for coordinating metabolism at the population level. This differentiation in response to environmental cues has been demonstrated in *E. coli* biofilms, where the amino acid L-alanine acts as a metabolic valve. L-alanine is secreted and utilized by a sub-population exposed to oxygen when a suitable carbon source is absent [38]. However, a clear link between metabolism and AI-2 mediated quorum sensing is still missing *in vivo*.

It is widely accepted that different endogenous *Enterobacteriaceae* offer different levels of protection against invading pathogens, such as *S*. Tm [39]. One of the deciding factors is metabolic resource overlap between the host microbiota, *E. coli*, and *S.* Tm that defines if a host is susceptible towards infection [40]. Since both species have a high metabolic resource overlap, we wondered what roles chemotaxis and quorum sensing play in enterobacterial competition.

In this study, we investigated the role of AI-2 mediated quorum sensing in interspecies competition between *E. coli* and *S.* Tm. Using competitive infections in a streptomycin-pretreated mouse model, we demonstrated that *E. coli* utilizes AI-2 chemotaxis to compete against *S.* Tm. Conversely, *S.* Tm requires chemotaxis, albeit not towards AI-2, to efficiently compete against resident *E. coli* and cause enterocolitis. We further explored the effect of *E. coli* AI-2-dependent niche colonization on central metabolism of *S.* Tm cells. For this, we utilized a previously published *S.* Tm mutant pool that probes key aspects of carbohydrate utilization and expanded its mutant range to include mutants involved in glycolytic pathways and mixed acid fermentation. We assessed how *E. coli*, both wild-type and AI-2 QS-deficient, influenced the fitness of these *S.* Tm mutants. Here, we present initial evidence showing how quorum sensing governs interspecies competition.

## Results

### AI-2 chemotaxis of *E. coli* is involved in *E. coli*-*S.* Tm competition *in vivo*

In our previous study, we demonstrated that LsrB-mediated chemotaxis towards the self-produced quorum sensing molecule AI-2 provides *E. coli* with a fitness advantage during mouse gut colonization [18]. We were therefore interested in whether such AI-2 chemotaxis-dependent gut colonization results in increased colonization resistance of *E. coli* against the closely related species, the enteric pathogen *S.* Tm. To test this hypothesis, we used the experimental setup shown in Fig 1a. Specific pathogen-free (SPF) C57BL/6 mice were orally treated with either streptomycin or ampicillin to break colonization resistance and allow *E. coli* and *S.* Tm colonization [41,42]. One day post antibiotic treatment, the mice were orally infected with *E. coli* Z1331 (human commensal isolate) wild-type or Δ*lsrB* (no chemotaxis towards AI-2) [18,43,44], followed by challenge with *S.* Tm SL1344 wild-type strain 24 h later. The CFUs of both species were monitored daily for the next 4 days. On day 4 post infection (p.i.), mice were euthanized, and systemic spread of *S.* Tm was analyzed by collecting and plating mesenteric lymph nodes (mLNs), liver and spleen. As shown in Fig 1b and Fig S1a-b, although *S.* Tm is capable of growth in *E. coli*-precolonized mice, the *S.* Tm-*E. coli* CFU ratio is significantly higher in *E. coli* Δ*lsrB*-precolonized mice compared to the *E. coli* wild-type group. This indicated that AI-2 chemotaxis indeed contributes to *E. coli*-*S.* Tm competition *in vivo*. In agreement with these observations, similar results were obtained in mice that were precolonized with a non-chemotactic *E. coli* Δ*cheY* strain (Fig S1c). The presence of neither *E. coli* WT, Δ*lsrB* nor Δ*cheY* significantly affected systemic CFU loads of *S.* Tm (Fig S1d).

**Fig. 1.**
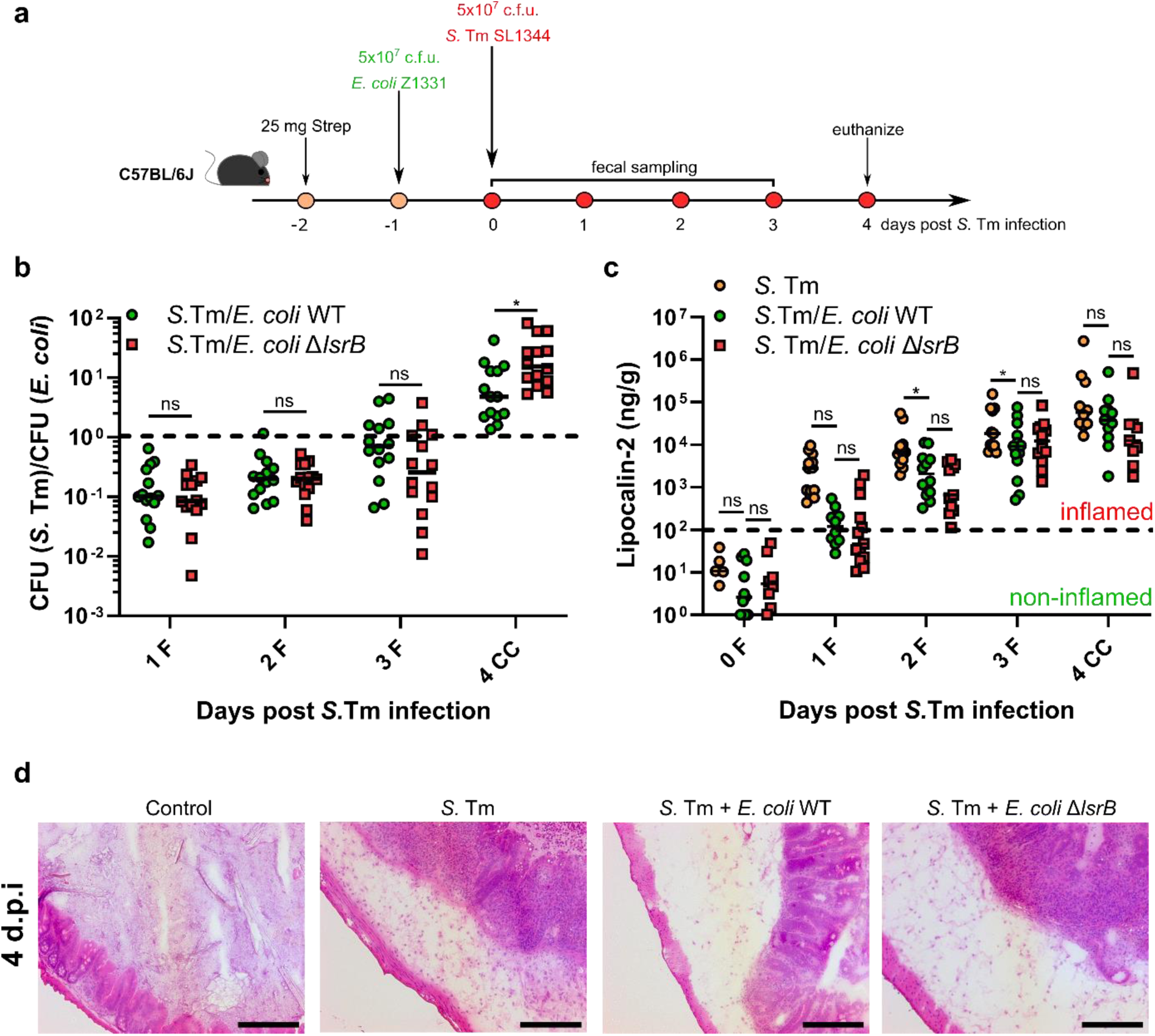
AI-2 chemotaxis-dependent *S.* Tm-*E. coli* competition in the murine gut. **(a)** Experimental scheme of competitive infections. C57BL/6J specific pathogen-free mice were pretreated with 25 mg of streptomycin and pre-colonized with *E. coli* Z1331 by oral gavage, followed by oral *S.* Tm infection. Feces were collected at 0, 1, 2, 3 days post *S.* Tm infection, and mice were euthanized at 4 d.p.i. **(b)** Competitive infections of *S.* Tm SL1344 against resident *E. coli* Z1331 wild-type or AI-2 chemotaxis-negative Δ*lsrB* mutant strain. The lines indicate median values (mice n=14, at least two independent experiments). P values were calculated using the two-tailed Mann-Whitney *U*-test (*P<0.05, ns – not significant). The dashed line indicates the CI value of 1. F, feces; CC, cecal content. **(c)** Lipocalin-2 levels in (F) feces and (CC) cecal content of mice infected with *S.* Tm, with or without pre-colonization with *E. coli* wild-type or Δ*lsrB* strains. Dashed line indicates approximate level of lipocalin-2 marking a shift towards gut inflammation. Lines indicate median values (min mice n=4, at least two independent experiments). P values were calculated using the two-tailed Mann-Whitney *U*-test (*P<0.05, ns – not significant). **(d)** Representative H&E staining images of cecal tissue of *S.* Tm-infected mice at 4 d.p.i. Mice infected with avirulent *S.* Tm SL1344 Δ*invG* Δ*sseD* strain were used as a control. Scale bars, 200 μm.

In line with delayed *S*. Tm gut colonization kinetics (Fig S1a), less gut inflammation was observed in *E. coli*-colonized mice for the first 2 days of *S.* Tm infection, as measured by lipocalin-2 levels (Fig 1c). The levels of inflammation evened out at days 3 and 4 p.i., with no significant differences between the *E. coli* groups. Consistent with the lipocalin-2 data, the analysis of the cecal tissue pathology at 4 d.p.i. revealed similar levels of pathological changes (Fig 1d). Our findings thus suggest that AI-2 chemotaxis plays a role in *E. coli*-*S.* Tm competition, albeit without affecting the capability of *S.* Tm to spread systemically.

### Chemotaxis provides *S.* Tm with a fitness advantage in *E. coli*-precolonized mice

Similarly to *E. coli*, *S.* Tm SL1344 benefits from chemotaxis during gut infection, as previously shown in a competitive infection model [13,45]. However, in contrast to *E. coli*, the fitness advantage of chemotaxis in *S.* Tm only becomes apparent with the onset of gut inflammation, and no fitness defect was observed for *S.* Tm Δ*cheY* mutant in its absence [14]. Having shown that AI-2 chemotaxis enhances colonization resistance of *E. coli* against *S.* Tm, we next hypothesized that *S.* Tm might employ the same strategy to invade the mouse gut precolonized with *E. coli*. Importantly, *S.* Tm SL1344 also encodes a functional *lsr* operon, potentially allowing it to profit from AI-2 chemotaxis during infection [46,47].

To test this, we used the experimental setup described above, competing *S.* Tm SL1344 wild type, Δ*cheY* and Δ*lsrB* strains against *E. coli* Z1331. The non-chemotactic *S.* Tm SL1344 Δ*cheY* strain showed a pronounced fitness defect in *E. coli*-precolonized mice, as evidenced by the decreased *S.* Tm-*E. coli* CFU ratio (Fig 2a). Additionally, the Δ*cheY* mutant caused significantly less gut inflammation and pathology during the first 2 days of infection (Fig 2b-d). On the other hand, no colonization defect or changes in gut inflammation levels were observed for *S.* Tm Δ*lsrB* throughout the course of the experiment (Fig 2a-d). Collectively, our observations suggest that, although chemotaxis enhances *S.* Tm gut colonization and the development of enterocolitis in the presence of *E. coli*, chemotactic cues other than AI-2 are involved in this process.

**Fig 2.**
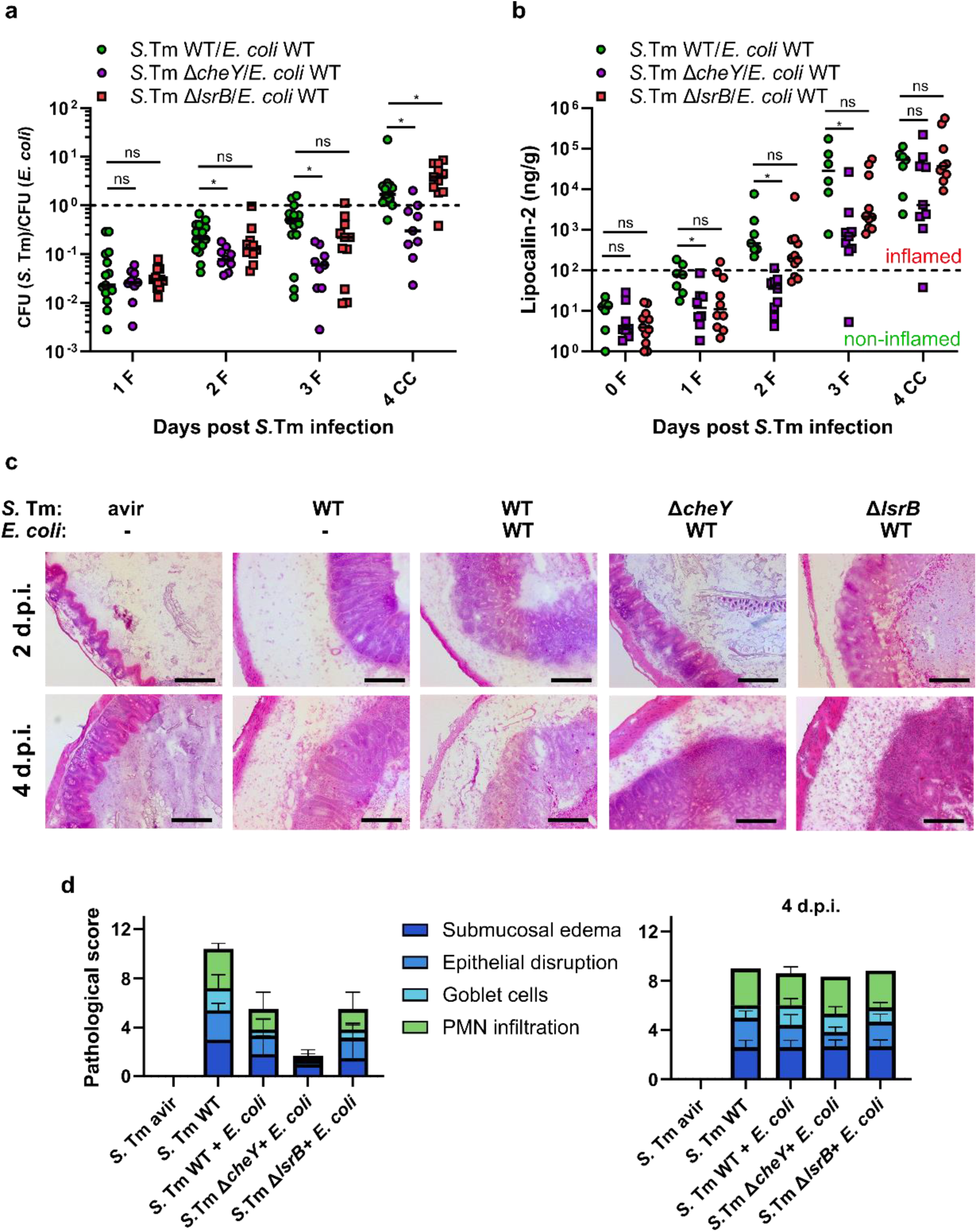
*S.* Tm requires chemotaxis, albeit not towards AI-2, to compete against resident *E. coli* during gut infection. **(a)** Competitive infection of *S.* Tm SL1344 wild-type and its non-chemotactic Δ*cheY* or non-AI-2-chemotactic Δ*lsrB* mutant strains against resident *E. coli* Z1331 strain. The lines indicate median values (min mice n=9, at least two independent experiments). P values were calculated using the two-tailed Mann-Whitney *U*-test (*P<0.05, ns – not significant). The dashed line indicates the CI value of 1. F, feces; CC, cecal content. **(b)** Lipocalin-2 levels in (F) feces and (CC) cecal content of mice infected with *S.* Tm as described in panel (a). Dashed line indicates approximate level of lipocalin-2 marking a shift towards gut inflammation. Lines indicate median values (min mice n=7, at least two independent experiments). P values were calculated using the two-tailed Mann-Whitney *U*-test (*P<0.05, ns – not significant). **(c)** Representative H&E staining images of cecal tissue of *S.* Tm-infected mice at 2 and 4 d.p.i. Scale bars, 200 μm. **(d)** Histopathology analysis of the cecal tissue as seen above in panel (c). Sections from at least four mice per group were analyzed.

To further dissect the role of chemotaxis in *S.* Tm infection, we analyzed the fitness of *S*. Tm SL1344 Δ*cheY* knockout strain relative to the wild-type strain in mice that were either precolonized with *E. coli* or not. In the competitive infection without *E. coli*, we observed a progressive decrease of the competitive index (CI) values between the mutant and the wild-type strain, indicating a strong fitness disadvantage for the non-chemotactic strain (Fig 3a, Fig S2a). Somewhat counterintuitively, the presence of *E. coli*, a close relative and thus a likely competitor of *S.* Tm [33,39,48], partially ameliorated the fitness disadvantage of *S.* Tm Δ*cheY* strain compared to the wild type. Knowing that the advantage of chemotaxis in *S.* Tm SL1344 is linked to inflammation, we reasoned that its influence on the outcome of *S.* Tm-*E. coli* competition is higher than the mere presence of *E. coli*. This hypothesis is supported by our previous observations, which show that the levels of *S.* Tm-induced inflammation are indeed lower in mice precolonized with *E. coli* (Fig 1c). Furthermore, in an avirulent *S.* Tm strain background (Δ*invG* Δ*sseD,* [49]) which is incapable of tissue invasion and induction of inflammation no fitness disadvantage of *S.* Tm Δ*cheY* strain was observed, regardless of the presence of *E. coli* (Fig 3b, Fig S2b-c). Consistent with these observations, non-chemotactic *S.* Tm Δ*cheY* did not show a growth defect in the gut lumen of *E. coli*-precolonized mice in absence of inflammation (Fig S3).

**Fig 3.**
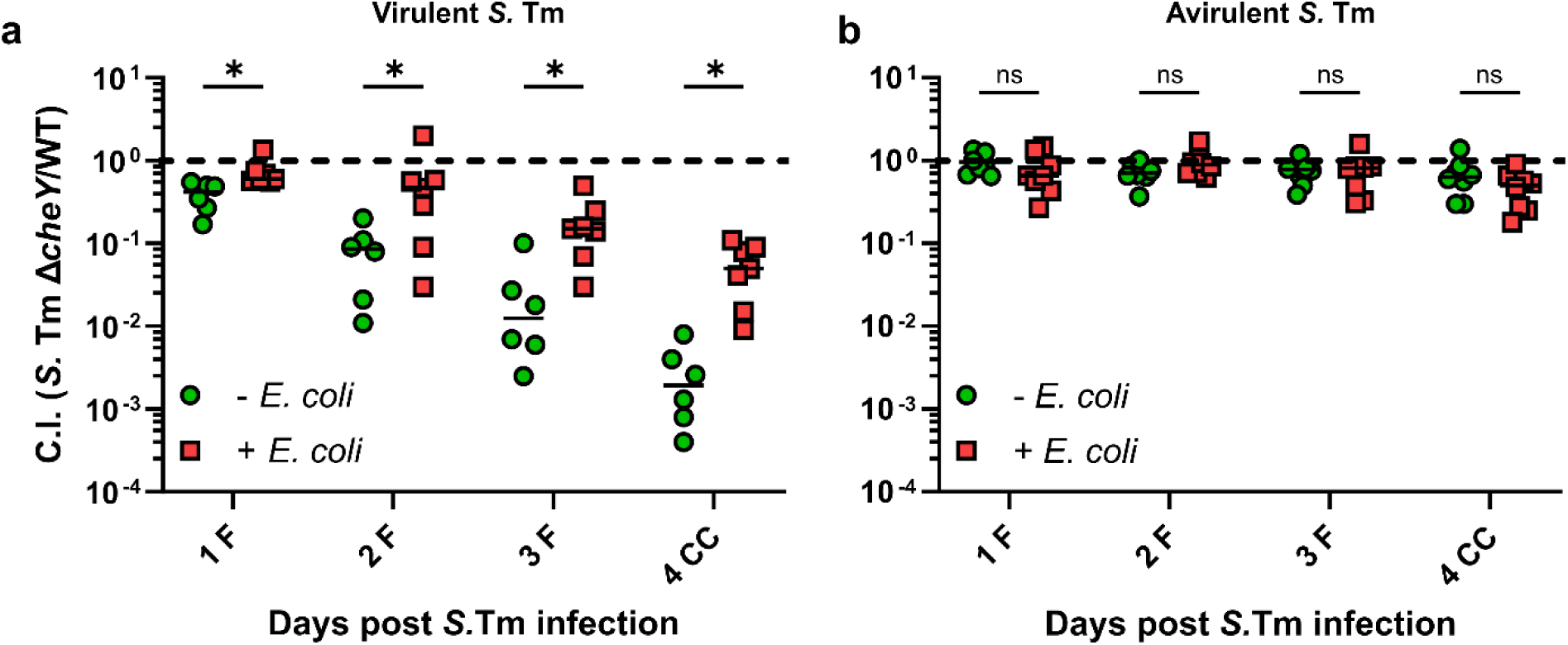
The relative fitness of *S.* Tm Δ*cheY* knockout strain is indirectly influenced by resident *E. coli* via delay of inflammation. Competitive infection *S.* Tm SL1344 Δ*cheY* knockout strain against the wild-type strain in **(a)** wild-type virulent or **(b)** avirulent Δ*invG* Δ*sseD* background. The lines indicate median values (min mice n=6, at least two independent experiments). P values were calculated using the two-tailed Mann-Whitney *U*-test (*P<0.05, ns – not significant). The dashed line indicates the CI value of 1. F, feces; CC, cecal content.

### *E. coli* presence in gut lumen leads to altered metabolic requirements for intraluminal growth of *S.* Tm

Our previous study showed that AI-2 chemotaxis, in addition to enhancing gut colonization by *E. coli*, leads to niche segregation of different *E. coli* strains based on their ability to sense AI-2, resulting in co-existence of such strains [18]. This implies that *E. coli*, based on its ability to sense and to respond chemotactically to AI-2, may occupy discrete niches within the gut, potentially altering the metabolite profile available to the invading *Salmonella* strain.

To investigate how AI-2-mediated quorum sensing influences the metabolic requirements for the intraluminal growth of *S*. Tm, we employed a WISH-barcoded carbohydrate utilization mutant pool [50].The *E. coli* non AI-2 chemotactic Δ*lsrB* mutant is impaired in AI-2 transport [37,43,51], thereby affecting AI-2-mediated transcriptional and post-translational regulation and chemotaxis towards AI-2 [44,46,51]. It allowed us to explore the role of AI-2-mediated quorum sensing in the competition between *S*. Tm and *E. coli* in context of metabolic exploitation. The experiment followed the same experimental scheme as described above (Fig 1a). Importantly, utilization of the WISH-barcoded *S*. Tm pool did not alter the overall *S.* Tm-*E. coli* competition outcome (compare Fig 1b and Fig S4a)

The *S*. Tm pool included seven wild-type strains to evaluate population bottlenecks during colonization, coinciding with the onset of inflammation. On days 1 and 2 post infection, the Shannon Evenness Score (SES) was close to 1 in all mice, indicating that we can reliably interpret the mutant fitness defects in the animals. By day 3, it dropped noticeably (Fig S4b). Consistent with the lipocalin-2 data shown in Fig 1c, we observed a tendency for *E. coli* to mitigate *S*. Tm-mediated inflammation, evident from the overall higher SES (Fig S4b). The interpretation of the CIs was limited to 2 days post-infection, as a significant number of samples fell below the 0.9 SES threshold for days 3 and 4 post infection, previously established as a cutoff [50,52]. Additionally, three control mutants (Δ*dcuABC*, Δ*frd*, and Δ*hyb*), targeting fumarate respiration and hydrogen utilization, showed similar CIs as previously [50,53], verifying the data (Fig 4a-b, Fig S5).

**Fig. 4.**
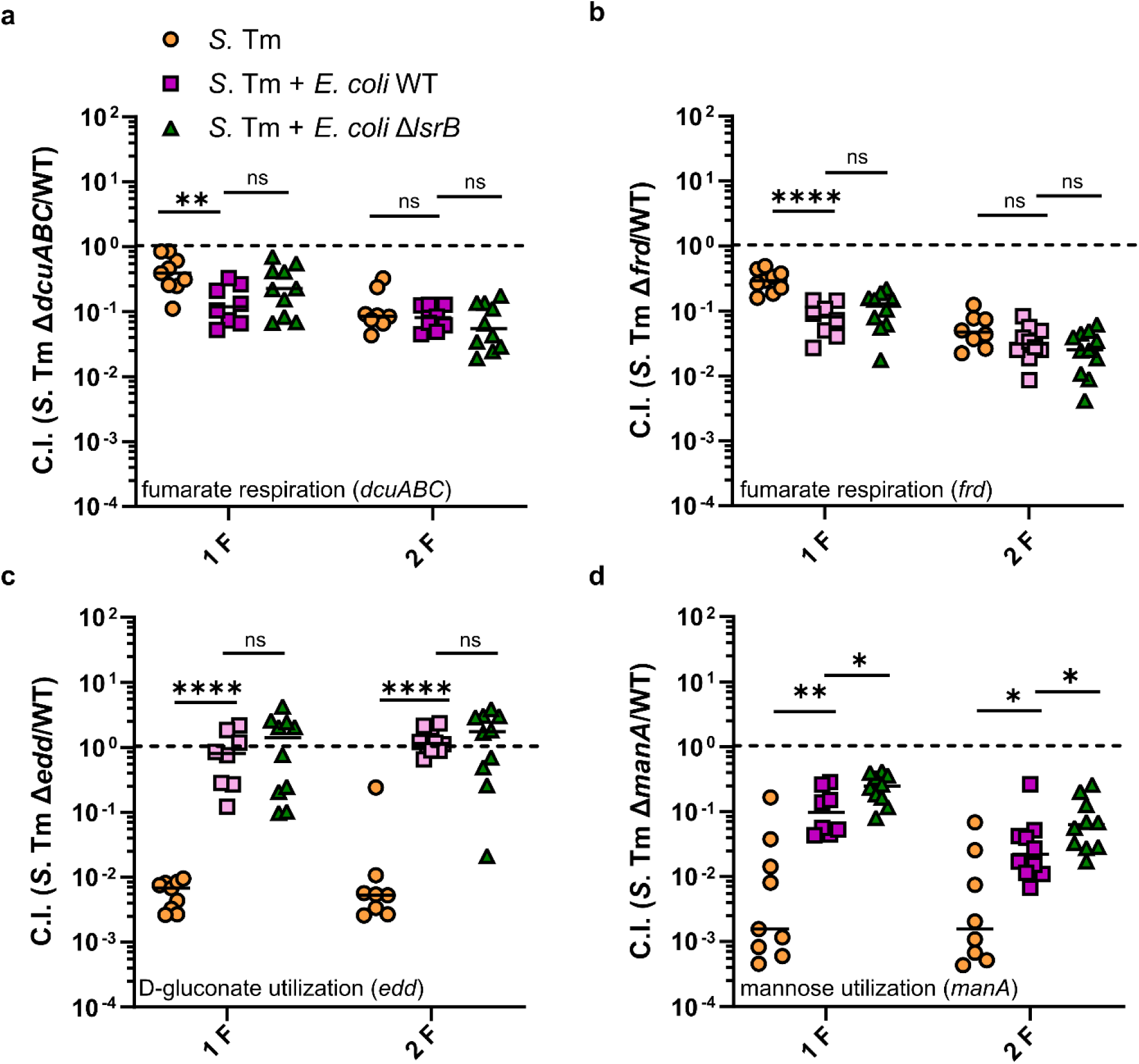
Resident *E. coli* affects *S.* Tm central carbon metabolism during gut infection in both AI-2-dependent and -independent manners. CI values of *S.* Tm mutants lacking **(a)** C4-dicarboxylate antiporters (Δ*dcuABC*), **(b)** terminal fumarate reductase (Δ*frd*), **(c)** phopshogluconate dehydratase (Δ*edd*) and **(d)** mannose-6-phophate isomerase (Δ*manA*), respectively. The plotted data represents the CI values calculated for the *S.* Tm mutant pool (see Fig. S5). The lines indicate median values (min mice n=8, at least two independent experiments). P values were calculated using the two-tailed Mann-Whitney *U*-test (****P<0.0001, **P<0.005, *P<0.05, ns – not significant). The dashed line indicates the CI value of 1. F, feces; CC, cecal content.

The pool included metabolic mutants deficient in carbohydrate utilization, mutants of the three glycolytic pathways, and mixed acid fermentation (Fig S5). A comprehensive list of these mutants and the CI values is provided in Tables S1 and S4.

The analysis of the barcoded mutants indicated that the presence of *E. coli* did not seem to have a profound effect on *S.* Tm metabolism, with only several *S.* Tm mutants being affected during the first 2 days of infection (Fig S5). The most prominent example of such a change was observed for the Δ*edd* mutant. The *edd* gene encodes phosphogluconate dehydratase, a key enzyme for utilizing the sugar acid D-gluconate via the Entner-Doudoroff pathway. While being disadvantageous in *S.* Tm single infection, the Δ*edd* mutation showed no loss of fitness in the gut that was precolonized with either the wild-type *E. coli* or Δ*lsrB* strain (Fig 4c).

The *dcuA*, *dcuB*, and *dcuC* genes (Δ*dcuABC*) encode C4-dicarboxylate antiporters, while *frd* operon encodes the terminal fumarate reductase. These genes have been previously demonstrated to play a significant role in the colonization of *S*. Tm and *E. coli* [53–57]. The presence of *E. coli* wild-type and Δ*lsrB* had similar effects on *S*. Tm Δ*dcuABC* and Δ*frd* mutants, significantly reducing their fitness at 1 day post infection (Fig 4a-b). However, this effect was lost by day 2 post *S.* Tm infection. Fumarate respiration has been identified as a critical factor in the competitive interactions between *S*. Tm and *E. coli* during colonization [53,54]. For this reason, a follow-up competitive 1:1 infection study was conducted involving an *S*. Tm wild-type and Δ*dcuABC* knockout strain in the presence of *E. coli* or its Δ*lsrB* mutant, following the experimental scheme shown in Fig 1a. This allowed us to track the fitness of the mutant strain for all 4 days of infection. As expected, no difference in the fitness of the *S.* Tm Δ*dcuABC* mutant was detected between the *E. coli* wild-type and Δ*lsrB* groups during the first 2 days of infection (Fig. 5a). However, at 3 days post infection, we observed significant loss of fitness of the mutant strain in mice precolonized with *E. coli* Δ*lsrB* as compared to those precolonized with wild-type *E. coli*. The same tendency, although statistically insignificant, was observed at 4 days post infection as well. This suggests that AI-2-dependent niche occupation by *E. coli* relieves the pressure on *S.* Tm to utilize fumarate respiration at later stages of infection.

**Fig. 5.**
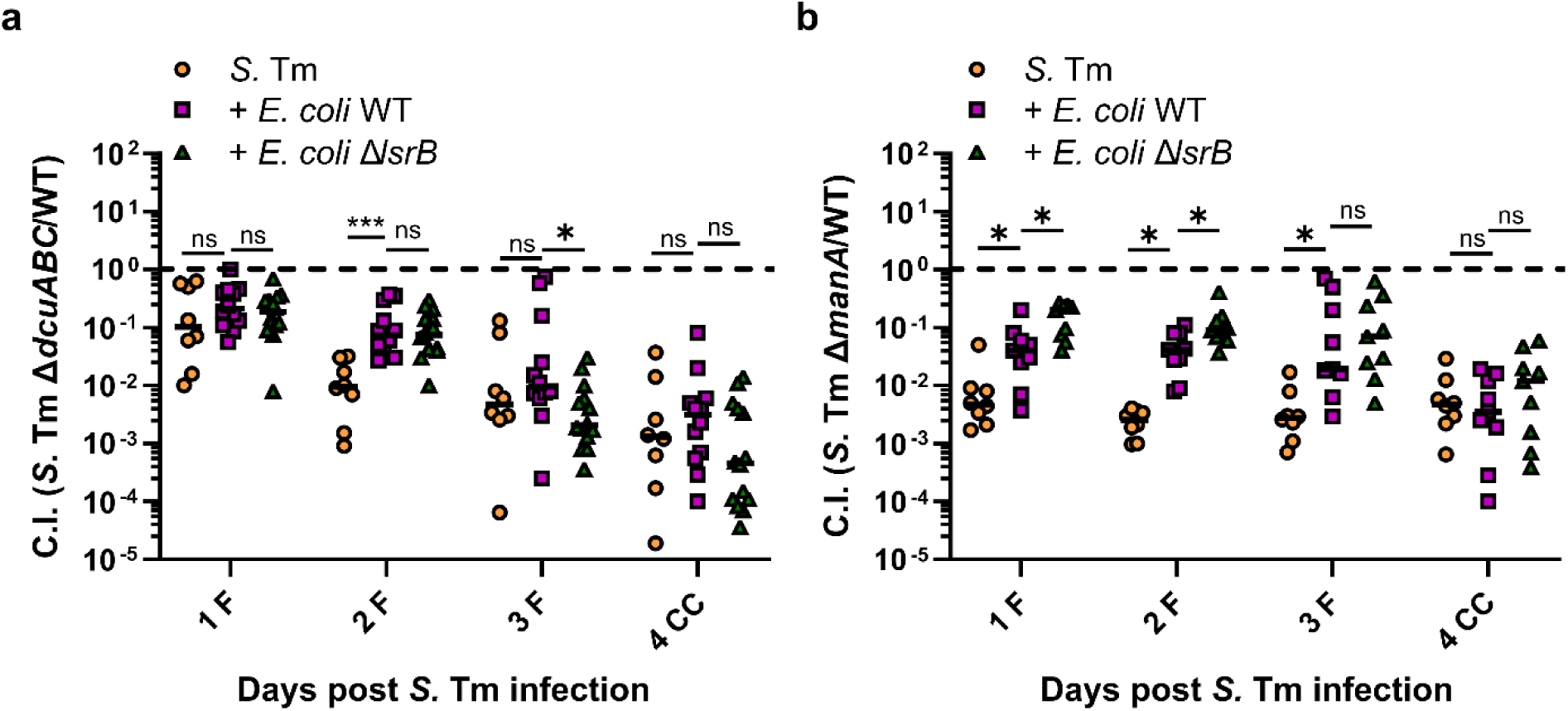
AI-2-dependent niche occupation by *E. coli* alters *S*. Tm metabolism *in vivo*. Competitive infections of S. Tm **(a)** - deficient Δ*dcuABC* and **(b)** mannose utilization-deficient Δ*manA* mutants against the wild-type strain in absence of *E. coli* as well as in presence of *E. coli* Z1331 wild-type or Δ*lsrB* strain. The lines indicate median values (min mice n=8, at least two independent experiments). P values were calculated using the two-tailed Mann-Whitney *U*-test (***P<0.0001, *P<0.05, ns – not significant). The dashed line indicates the CI value of 1. F, feces; CC, cecal content.

Finally, the most pronounced AI-2-dependent differences were observed for the *S*. Tm Δ*manA* mutant, which lacks mannose-6-phosphate isomerase, an enzyme crucial for the utilization of D-mannose. Similarly to Δ*dcuABC*, the presence of *E. coli* seemed to abolish the fitness disadvantage of Δ*manA* mutant that was observed in mice infected only with *S.* Tm (Fig 4d, Fig 5b). However, contrary to Δ*dcuABC*, intact AI-2 signaling in *E. coli* negatively affected the fitness of *S.* Tm Δ*manA*, and this effect was observed only during the first 2 days of *S.* Tm infection. At 3 days post infection, although still showing less fitness disadvantage compared to mice infected only with *S.* Tm, no difference in the fitness of *S.* Tm Δ*manA* was observed between mice colonized with *E. coli* wild-type or Δ*lsrB* strain. All groups showed a similar loss of fitness in the mannose utilization-deficient *S.* Tm strain by 4 days post infection. Collectively, our data provides evidence that AI-2-dependent gut colonization by *E. coli* has specific effects on the fitness profiles of mutants with defects in pathways for central metabolism of the invading *S.* Tm strain.

## Discussion

Freter’s nutrient niche theory posits that a bacterium can only colonize the gut if it can most efficiently utilize at least one specific limiting nutrient [58–60]. This theory can be expanded into the Restaurant Hypothesis by incorporating the concepts of spatial competition and the highly heterogeneous environment of the gut [61,62]. Investing energy into motility and chemotaxis, by allowing bacteria to efficiently navigate the chemical gradients in the gut, represents an important survival strategy compared to a non-motile lifestyle [4]. The growing body of work clearly shows the importance of chemotaxis in environmental and host-associated bacteria [4,5]. However, as the gut ecosystem is characterized by a very complex and heterogeneous chemical environment, it is not always feasible to identify specific chemoattractants or chemorepellents that contribute to gut colonization [63].

The connection between chemotaxis and quorum sensing in *E. coli* has been established in previous studies [44], showing that chemotaxis towards AI-2 enhances gut colonization and can result in niche segregation between AI-2-chemotactic and non-AI-2-chemotactic *E. coli* strains [18]. Interestingly, the apparent effect of AI-2 chemotaxis was linked to ability of *E. coli* to consume fructoselysine, establishing a further connection between central metabolism and collective behavior via the quorum sensing system. However, it is not well understood how AI-2-mediated quorum sensing is involved in intraspecies competition *in vivo*, such as between *S.* Tm and *E. coli*.

In this work, we show that AI-2-dependent gut colonization by *E. coli* Z1331 results in higher colonization resistance against invading *S.* Tm SL1344, as both non-chemotactic and AI-2 chemotaxis-deficient mutants seem to compete less efficiently against *S*. Tm at 4 days post infection. Since *S.* Tm is a close relative of *E. coli*, we assumed that AI-2 chemotaxis might contribute to its competition against the resident *E. coli* strain as well. Like *E. coli*, *S.* Tm possesses a functional *lsr* operon [46]. In our experiments, although the non-chemotactic *S.* Tm mutant clearly showed a competitive disadvantage, resulting in slower kinetics of gut inflammation, no such defects were observed for the AI-2 chemotaxis-deficient mutant. These observations underscore that even closely related species might employ different chemotactic cues to locate their respective niches within the gut. Accordingly, *S.* Tm strains possess a slightly different set of chemoreceptors, including some not found in *E. coli* [64].

In agreement with previous observations [13,14], the development of gut inflammation appeared to be the deciding factor for the fitness of the non-chemotactic *S.* Tm Δ*cheY* mutant. On the contrary, the presence of *E. coli* only seemed to indirectly affect the competitive phenotype of the *S.* Tm Δ*cheY* by further slowing down the onset of inflammation. A similar observation was reported with the murine isolate *E. coli* 8178, which reduced intestinal inflammation caused by *S.* Tm [39,53,65]. In the absence of inflammation, the chemotaxis system of *S.* Tm was dispensable for both intra- and interspecies competition *in vivo*. This further highlights a fundamental difference between the physiological roles of chemotaxis in *S.* Tm and *E. coli*. In the latter, the fitness advantage of chemotaxis is not dependent on inflammation and becomes apparent only in competitive infections [18,66].

However, and most intriguingly, the presence of *E. coli*, as well as its ability to occupy its respective niche in an AI-2-dependent manner, appeared to affect the central metabolism of the invading *S.* Tm strain. Utilizing the recently developed library of WISH-barcoded carbohydrate utilization mutants of *S*. Tm SL1344 [50], we investigated fitness changes caused by an altered gut environment in an AI-2-dependent manner due to *E. coli*. Out of 49 tested mutants, altered fitness in the presence of *E. coli* during the first 2 days of infection, albeit not dependent on AI-2 signaling, was observed for the *S*. Tm Δ*frd* (fumarate respiration) and Δ*edd* (D-gluconate utilization) strains. Fumarate respiration has previously been reported to play a crucial role during initial growth [53,54], and late-stage infection, when inflammation is pronounced [54]. Therefore, it was not surprising that the presence of *E. coli*, which also relies on fumarate respiration [56], further decreased the fitness of the *S.* Tm Δ*frd* mutant. Contrarily, the *S.* Tm Δ*edd* mutant, significantly impaired in gut colonization in the absence of *E. coli*, showed neutral fitness in its presence.

Finally, we identified two metabolic pathways in *S.* Tm that were affected by AI-2 signaling in *E. coli*. One of them is fumarate respiration, already mentioned above. The fitness of the *S.* Tm Δ*dcuABC* mutant, lacking genes for C4-dicarboxylate antiporters and thus, like Δ*frd*, deficient in fumarate respiration, was positively affected by AI-2 signaling in *E. coli*. This was, however, only observed on day 3 of infection. During the first two days of *S.* Tm infection, we observed an opposite effect of *E. coli* AI-2 signaling on the fitness of the *S.* Tm mannose utilization-deficient Δ*manA* strain. AI-2-dependent niche occupation by *E. coli* resulted in increased pressure for *S.* Tm to metabolize mannose, resulting in lower fitness of the *S.* Tm Δ*manA* strain in mice precolonized with wild-type *E. coli* compared to those precolonized with the *E. coli* Δ*lsrB* strain. It is noteworthy that metabolic mutants, such as Δ*edd*, can have pleiotropic effects on sugar phosphate accumulation, which is not the case for *manA* [67,68]. In our previous study, we observed that the utilization of fructoselysine, which was either partially imported by the phosphotransferase system (PTS) or activated its components, resulted in more AI-2 production by *E. coli* by inhibiting *lsr* operon activity [18]. Increased AI-2 production by the fructoselysine-consuming population of *E. coli* might thus attract further *E. coli* cells to the source of fructoselysine in the gut by means of AI-2 chemotaxis. As mannose is also imported via the PTS system [69], one could speculate that a similar mechanism is in place. This might result in more efficient consumption of mannose by wild-type *E. coli* compared to AI-2 chemotaxis-deficient *E. coli* Δ*lsrB* and explain the observed changes in the fitness of the *S*. Tm Δ*manA* mutant. However, further studies are needed to explain this observation.

Admittedly, a clear mechanistic understanding of how quorum sensing-dependent niche colonization by *E. coli* affects its interspecies interactions is still missing and requires more studies. However, this work highlights the fact that the AI-2 quorum sensing signaling of one species might affect the metabolic environment *in vivo* and, by proxy, the metabolism of its interaction partners in complex communities. Quorum quenching involves the inhibition of quorum sensing (QS) through chemicals or enzymes, effectively preventing bacterial communication [70,71]. Considering these presented results, this approach could lead to new strategies for specifically altering the metabolic environment of the gut lumen through a probiotic. In combination with quorum quenching, this could open up the possibility of inhibiting pathogen communication and preventing population-wide adaptation, ultimately limiting the ability of pathogens to colonize the mammalian gut.

## Materials and Methods

### Bacterial strains and growth conditions

The *E. coli* and *S.* Tm strains, plasmids and oligos used in this study are listed in Tables S4-6. The strains were routinely cultivated in liquid Lysogeny Broth (LB) or on 1.5% LB agar supplemented with kanamycin (50 µg/ml), ampicillin (100 µg/ml), streptomycin (50 µg/ml) or chloramphenicol (35 µg/ml), where necessary. Gene knockouts or chromosomal integrations were obtained via lambda-red recombination [72] and P22 transduction. Streptomycin resistance, conferred by the *S.* Tm SL1344 P3 plasmid, was introduced into *E. coli* strains via conjugation as previously described [73].

### Homologous recombination by Lambda Red

Single-gene knockout strains were generated using the lambda-red single-step protocol [72]. Primers were designed with an approximately 40 bp overhanging region homologous to the genomic region of interest and 20 bp binding region corresponding to the antibiotic resistance cassette (Table S6). PCR amplification was performed using the plasmid pKD4 for kanamycin resistance or the pTWIST plasmids for WISH tags, which include an ampicillin resistance cassette. DreamTaq Master Mix (Thermo Fisher Scientific) was employed, followed by digestion of the template DNA using FastDigest DpnI (Thermo Fisher Scientific). Subsequently, the PCR product was purified using the Qiagen DNA purification kit. SB300 with either the pKD46 or pSIM5 plasmid was cultured for 3 h at 30°C until early exponential phase, followed by induction with L-arabinose (10 mM, Sigma-Aldrich) or 42°C for 20 min, respectively. The cells were washed in ice-cold glycerol (10% v/v) solution and concentrated 100-fold. Ultimately, the PCR product was transformed by electroshock (1.8 V at 5 ms), followed by regeneration in SOC (SOB pre-made mixture, Roth GmbH, and 50 mM glucose) medium for 2 h at 37°C, ultimately plated on selective LB-agar plates. The success of the gene knockout was verified by gel electrophoresis and sanger sequencing (Microsynth AG). Kanamycin resistance cassettes were eliminated via flippase FLP recombination [74].

### Homologous recombination by P22 phage transduction

P22 phage transduction was conducted by generating P22 phages containing the antibiotic resistance cassette inserted into the gene of interest from the defined single-gene deletion mutant collection of *S.* Tm [75]. The single-gene knockout mutant was incubated overnight with the P22 phage generated from a wild type SB300 background. The culture was treated with chloroform (1% v/v) for 15 min followed by centrifugation and subsequent sterile filtration (0.44 µm pore size). The P22 phages were subsequently incubated with the recipient strain for 15 minutes and then plated on selective LB-agar plates. This was followed by two consecutive overnight streaks on selective LB-agar plates. Finally, the transduced clone was examined for P22 phage contamination using Evans Blue Uranine (EBU) LB-agar plates (0.4% w/v glucose, 0.001% w/v Evans Blue, 0.002% w/v Uranine). All mutations were verified by gel electrophoresis or Sanger sequencing (Microsynth AG), using the corresponding primers (Table S6).

### WISH-barcoding of S. Tm

WISH barcodes were introduced, as previously described [76]. WISH-tags were amplified from pTWIST using DreamTaq Master Mix (Thermo Fisher Scientific) with WISH_int_fwd and WISH_int_rev primers (Table S6) and integrated into *S.* Tm SL1344 (strain SB300), using the λ-red system with pSIM5 [77]. Integration was targeted at a fitness-neutral locus between the pseudogenes *malX* and *malY*, as previously described [78]. Correct integration was confirmed through colony PCR, and WISH-tags were validated by Sanger sequencing (Microsynth AG), using either the WISH_ver_fwd and WISH_ver_rev primers or the WISH_seq_fwd and WISH_seq_rev primers (Table S6). Subsequently, P22 phage lysates were prepared from these generated strains to transduce the WISH-tag into *S.* Tm SL1344 (SB300) mutants of the three glycolytic pathways and mixed acid fermentation.

### Animals

C57BL/6 (JAX:000664, The Jackson Laboratory) mice were used in all experiments. The mice were held under specific pathogen-free (SPF) conditions at the EPIC facility (ETH Zurich). The light/dark cycle was set to 12:12 h, with room temperature and humidity maintained at 21±1 °C and 50±10%, respectively. All experiments complied with cantonal and Swiss legislation and were approved by the Tierversuchskommission, Kantonales Veterinäramt Zürich under licenses ZH158/2019, ZH108/2022 and ZH109/2022.

### Mouse infection experiments

7–12-week-old mice SPF mice of both sexes were randomly assigned to experimental groups. The mice were orally pretreated with streptomycin (25 mg) or ampicillin (20 mg) 24 h prior to infection. *E. coli* and *S.* Tm cultures were grown overnight in LB at 37 °C with shaking, followed by dilution in 1:100 in fresh LB and incubation at 37 °C with shaking till the cultures reached mid-exponential phase of growth. The cells were then washed and resuspended in sterile PBS (137 mM NaCl, 2.7 mM KCl, 10 mM Na_2_HPO_4_, 1.8 mM KH_2_PO_4_). Unless stated otherwise, the mice were orally infected with 5×10^7^ CFU (in 50 µl) of *E. coli* or 50 µl PBS (as a control), followed by 5×10^7^ CFU (in 50 µl) *S.* Tm infection (wild-type or a 1:1 mixture of a wild-type and a knockout strain) 24 h post *E. coli* infection. Feces were collected every 24 h up to 4 days post S. Tm infection. At 2- or 4-days post *S.* Tm infection, mice were euthanized by CO_2_ asphyxiation. Cecal contents and systemic organs (mesenteric lymph nodes, liver (one sixth) and spleen) were harvested and suspended in 500 µl PBS, followed by homogenization in a Tissue Lyzer (Qiagen). Bacteria were plated on MacConkey (Oxoid) or LB agar plates with appropriated antibiotics to count *E. coli*, *S.* Tm wild-type and knockout cells.

Competitive index (CI) of a wild-type *S.* Tm strain and respective knockouts was determined as a ratio between the CFU counts of a knockout strain divided by that of the wild-type and normalized to the CI in the inoculum.

### Sample preparation for the WISH barcode counting

The mutant pool was prepared as previously described [50].Fecal *S*. Tm cells were enriched in 1 ml LB medium with 100 µg/ml carbenicillin (Carl Roth GmbH) to select for WISH-barcoded strains. Bacterial cells were pelleted, the supernatant was discarded, and then stored at −20 °C. DNA extraction from thawed pellets was performed using commercial kits (Qiagen Mini DNA kit) according to the manufacturer’s instructions. For PCR amplification of the WISH-barcodes, 2 µl of isolated genomic DNA sample and 100 nmol of each primer (WISH_Illumina_fwd and WISH_Illumina_rev, see Table S6) were used in a DreamTaq MasterMix (Thermo Fisher Scientific). The reaction was conducted with the following cycling program: initial denaturation step at (1) 95 °C for 3 min followed by (2) 95 °C for 30 sec, (3) 55 °C for 30 sec, (4) 72 °C for 20 sec for (5) 25 cycles, and a terminal extension step at (6) 72 °C for 10 min. PCR products were column purified. We indexed the PCR products for Illumina sequencing by performing a second PCR with nested unique dual index primers using the following program: (1) 95 °C for 3 min, (2) 95 °C for 30 s, (3) 55 °C for 30 s, (4) 72 °C for 20 sec, (5) repeat steps 2-4 for 10 cycles, (6) 72 °C for 3 min. Afterward, we assessed the indexed PCR product using gel electrophoresis (1% w/v agarose, TAE buffer), pooled the indexed samples according to band intensity, and subsequently purified the library via AMPure bead cleanup (Beckman Coulter) before proceeding to Illumina sequencing. Amplicon sequencing was performed by BMKGENE (Münster, Germany). BMKGENE was tasked with sequencing each sample at a 1 G output on the NGS Novaseq platform, utilizing a 150 bp paired end reads program. Subsequently, the reads were demultiplexed and grouped by WISH-tags using mBARq software [79]. Misreads or mutations of up to five bases were assigned to the closest correct WISH-tag sequence. The WISH barcode counts for each mouse in every experiment are available in Table S3. These counts were used to calculate the competitive fitness and Shannon evenness score (7 wild types). WISH counts with less than or equal to 10 were excluded from further analysis and defined as the detection limit, as previously defined [76]. As previously established [50], the competitive index of mutants that were below the limit of detection were conservatively set to a competitive index of 10^-3^.

### Competitive index calculation

To calculate the Competitive Index (CI) for the mutant pool, the values were determined by dividing the number of observed barcode reads at a specific time point (day 1 to day 4 post *S.* Tm infection or in cecum content) by the number of barcode reads observed in the inoculum, resulting in the individual strain fitness. For the calculation of the CI, the individual strain fitness of each WISH-barcoded mutant was divided by the mean fitness value of the 7 WISH-barcoded wild-type *S*. Tm control strains. To calculate the statistical significance, the metabolic mutants were compared to the SL1344 (SB300) wild type in the control group.

### Lipocalin-2 ELISA

Lipocalin-2 ELISA was used to analyze the levels of gut inflammation in the experiments. The measurements were performed on feces and cecal contents that had been previously homogenized in 500 μl PBS by ELISA DuoSet Lipocalin ELISA kit according to the manufacturer’s manual (DY1857, R&D Systems).

### Histopathology

Additionally to Lipocalin-2 ELISA, histopathology analysis was performed on cecal tissue sections to further analyse inflammatory response. Cecal tissue samples were embedded and snap-frozen in Tissue-Tek OCT medium (Sysmex), and 10 um cryosections were stained with haematoxylin and eosin (H&E). Pathological analysis and scoring (submucosal edema, numbers of goblet cells, epithelial integrity and polymorphonuclear granulocytes infiltration into lamina propria) was performed as previously described in blinded manner [42].

### Statistical analysis

The sample size in mouse experiments was not pre-determined, and mice of both sexes were randomly assigned to experimental groups. The two-tailed Mann Whitney-*U* test was used to compare the experimental groups, with *P* values of less than 0.05 indicating statistical significance. Statistical analysis was performed in GraphPad Prism v10 for Windows (GraphPad Software).

## Acknowledgements

The authors would like to acknowledge the staff at the ETH animal facilities (EPIC and RCHCI, especially Manuela Graf). This work has been funded by grants from the Swiss National Science Foundation (310030_192567, 10.001.588 and NCCR Microbiomes grant 51NF40_180575) to W.-D.H. C.S. is supported by the German Research Foundation (SCHU 3606/1-1).

## Author contributions

L.L., C.S. and W.-D.H conceived the study, L.L. and C.S. performed the experiments, M.B. prepared and stained cecal tissue, B.N. performed histopathology analysis, T.L. generated several *S.* Tm strains, C.S. and J.N. analyzed the WISH-barcoded mutant pool data. L.L. and C.S. wrote the manuscript, which was edited by all the authors.

## References

1. Hazelbauer GL. Bacterial chemotaxis: The early years of molecular studies. Annu Rev Microbiol. 2012;66: 285–303. doi:10.1146/annurev-micro-092611-150120

2. Kentner D, Sourjik V. Spatial organization of the bacterial chemotaxis system. Curr Opin Microbiol. 2006;9: 619–624. doi:10.1016/j.mib.2006.10.012

3. Yang Y, Pollard AM, Höfler C, Poschet G, Wirtz M, Hell R, et al. Relation between chemotaxis and consumption of amino acids in bacteria. Mol Microbiol. 2015;96: 1272–1282. doi:10.1111/mmi.13006

4. Keegstra JM, Carrara F, Stocker R. The ecological roles of bacterial chemotaxis. Nat Rev Microbiol. 2022;1–14. doi:10.1038/s41579-022-00709-w

5. Matilla MA, Krell T. The effect of bacterial chemotaxis on host infection and pathogenicity. FEMS Microbiol Rev. 2018;42: 40–67. doi:10.1093/femsre/fux052

6. Garvis S, Munder A, Ball G, De Bentzmann S, Wiehlmann L, Ewbank JJ, et al. *Caenorhabditis elegans* semi-automated liquid screen reveals a specialized role for the chemotaxis gene *cheB2* in *Pseudomonas aeruginosa* virulence. PLOS Pathog. 2009;5: e1000540. doi:10.1371/journal.ppat.1000540

7. Andermann TM, Chen Y-T, Ottemann KM. Two predicted chemoreceptors of *Helicobacter pylori* promote stomach infection. Infect Immun. 2002;70: 5877–5881. doi:10.1128/iai.70.10.5877-5881.2002

8. Huang JY, Goers Sweeney E, Guillemin K, Amieva MR. Multiple acid sensors control *Helicobacter pylori* colonization of the stomach. PLOS Pathog. 2017;13: e1006118. doi:10.1371/journal.ppat.1006118

9. Novak EA, Sekar P, Xu H, Moon KH, Manne A, Wooten RM, et al. The *Borrelia burgdorferi* CheY3 response regulator is essential for chemotaxis and completion of its natural infection cycle. Cell Microbiol. 2016;18: 1782–1799. doi:10.1111/cmi.12617

10. Xu H, Sultan S, Yerke A, Moon KH, Wooten RM, Motaleb MA. *Borrelia burgdorferi* CheY2 is dispensable for chemotaxis or motility but crucial for the infectious life cycle of the spirochete. Infect Immun. 2016;85(1): e00264–16. doi:10.1128/iai.00264-16

11. Sze CW, Zhang K, Kariu T, Pal U, Li C. *Borrelia burgdorferi* needs chemotaxis to establish infection in mammals and to accomplish its enzootic cycle. Infect Immun. 2012;80(7): 2485–2492. doi:10.1128/IAI.00145-12

12. Irazoki O, ter Beek J, Alvarez L, Mateus A, Colin R, Typas A, et al. D-amino acids signal a stress-dependent run-away response in *Vibrio cholerae*. Nat Microbiol. 2023;8: 1549–1560. doi:10.1038/s41564-023-01419-6

13. Stecher B, Hapfelmeier S, Müller C, Kremer M, Stallmach T, Hardt W-D. Flagella and chemotaxis are required for efficient induction of *Salmonella enterica* serovar Typhimurium colitis in streptomycin-pretreated mice. Infect Immun. 2004;72: 4138–4150. doi:10.1128/iai.72.7.4138-4150

14. Stecher B, Barthel M, Schlumberger MC, Haberli L, Rabsch W, Kremer M, et al. Motility allows *S.* Typhimurium to benefit from the mucosal defence. Cell Microbiol. 2008;10: 1166–1180. doi:10.1111/j.1462-5822.2008.01118.x

15. Mandel MJ, Schaefer AL, Brennan CA, Heath-Heckman EAC, DeLoney-Marino CR, McFall-Ngai MJ, et al. Squid-derived chitin oligosaccharides are a chemotactic signal during colonization by *Vibrio fischeri*. Appl Environ Microbiol. 2012;78: 4620. doi:10.1128/aem.00377-12

16. Graf J, Dunlap P V., Ruby EG. Effect of transposon-induced motility mutations on colonization of the host light organ by *Vibrio fischeri*. J Bacteriol. 1994;176: 6986–6991. doi:10.1128/jb.176.22.6986-6991.1994

17. Kajikawa A, Suzuki S, Igimi S. The impact of motility on the localization of *Lactobacillus agilis* in the murine gastrointestinal tract. BMC Microbiol. 2018;18. doi:10.1186/S12866-018-1219-3

18. Laganenka L, Lee J, Malfertheiner L, Dieterich C, Fuchs L, Piel J, et al. Chemotaxis and autoinducer-2 signalling enhance gut colonization and contribute to niche segregation of *Escherichia coli* strains in the mammalian gut. Nat Microbiol. 2023;8(2): 204–207. doi:10.1038/s41564-022-01286-7

19. Yang J, Chawla R, Rhee KY, Gupta R, Manson MD, Jayaraman A, et al. Biphasic chemotaxis of *Escherichia coli* to the microbiota metabolite indole. Proc Natl Acad Sci U S A. 2020;117: 6114–6120. doi:10.1073/pnas.1916974117

20. Croxen MA, Sisson G, Melano R, Hoffman PS. The *Helicobacter pylori* chemotaxis receptor TlpB (HP0103) is required for pH taxis and for colonization of the gastric mucosa. J Bacteriol. 2006;188: 2656–2665. doi:10.1128/jb.188.7.2656-2665.2006

21. Huang JY, Sweeney EG, Sigal M, Kuo CJ, Guillemin K, Amieva MR. Chemodetection and destruction of host urea allows *Helicobacter pylori* to locate the epithelium. Cell Host Microbe. 2015;18: 147–156. doi:10.1016/j.chom.2015.07.002

22. Selvaraj P, Gupta R, Peterson KM. The *Vibrio cholerae* ToxR regulon encodes host-specific chemotaxis proteins that function in intestinal colonization. SOJ Microbiol Infect Dis. 2015;3(3). doi:10.15226/sojmid/3/3/00141

23. Bansal T, Englert D, Lee J, Hegde M, Wood TK, Jayaraman A. Differential effects of epinephrine, norepinephrine, and indole on *Escherichia coli* O157:H7 chemotaxis, colonization, and gene expression. Infect Immun. 2007;75: 4597–4607. doi:10.1128/iai.00630-07

24. Pereira CS, Thompson J a., Xavier KB. AI-2-mediated signalling in bacteria. FEMS Microbiol Rev. 2013;37: 156–181. doi:10.1111/j.1574-6976.2012.00345.x

25. Ismail AS, Valastyan JS, Bassler BL. A host-produced autoinducer-2 mimic activates bacterial quorum sensing. Cell Host Microbe. 2016;19: 470–480. doi:10.1016/j.chom.2016.02.020

26. Thompson JA, Oliveira RA, Ubeda C, Xavier KB, Djukovic A. Manipulation of the quorum sensing signal AI-2 affects the antibiotic-treated gut microbiota. Cell Rep. 2015;10: 1861–1871. doi:10.1016/j.celrep.2015.02.049

27. Thompson JA, Oliveira RA, Xavier KB. Chemical conversations in the gut microbiota. Gut Microbes. 2016;7: 163–170. doi:10.1080/19490976.2016.1145374

28. Winter SE, Winter MG, Xavier MN, Thiennimitr P, Poon V, Keestra AM, et al. Host-derived nitrate boosts growth of *E. coli* in the inflamed gut. Science. 2013;339(6120):708–11. doi:10.1126/science.1232467

29. Rivera-Chávez F, Zhang LF, Faber F, Lopez CA, Byndloss MX, Olsan EE, et al. Depletion of butyrate-producing *Clostridia* from the gut microbiota drives an aerobic luminal expansion of *Salmonella*. Cell Host Microbe. 2016;19: 443–454. doi:10.1016/j.chom.2016.03.004

30. Winter SE, Thiennimitr P, Winter MG, Butler BP, Huseby DL, Crawford RW, et al. Gut inflammation provides a respiratory electron acceptor for *Salmonella*. Nature. 2010;467: 426–429. doi:10.1038/nature09415

31. Chang DE, Smalley DJ, Tucker DL, Leatham MP, Norris WE, Stevenson SJ, et al. Carbon nutrition of *Escherichia coli* in the mouse intestine. Proc Natl Acad Sci U S A. 2004;101: 7427– 7432. doi:10.1073/pnas.0307888101/

32. Fabich AJ, Jones SA, Chowdhury FZ, Cernosek A, Anderson A, Smalley D, et al. Comparison of carbon nutrition for pathogenic and commensal *Escherichia coli* strains in the mouse intestine. Infect Immun. 2008;76: 1143–1152. doi:10.1128/iai.01386-07

33. Eberl C, Weiss AS, Jochum LM, Durai Raj AC, Ring D, Hussain S, et al. *E. coli* enhance colonization resistance against *Salmonella* Typhimurium by competing for galactitol, a context-dependent limiting carbon source. Cell Host Microbe. 2021;29: 1680–1692.e7. doi:10.1016/j.chom.2021.09.004

34. Gül E, Abi Younes A, Huuskonen J, Diawara C, Nguyen BD, Maurer L, et al. Differences in carbon metabolic capacity fuel co-existence and plasmid transfer between *Salmonella* strains in the mouse gut. Cell Host Microbe. 2023;31: 1140–1153.e3. doi:10.1016/j.chom.2023.05.029

35. Ha J-H, Hauk P, Cho K, Eo Y, Ma X, Stephens K, et al. Evidence of link between quorum sensing and sugar metabolism in *Escherichia coli* revealed via cocrystal structures of LsrK and HPr. Sci Adv. 2018;4: eaar7063. doi:10.1126/sciadv.aar7063

36. Wang L, Hashimoto D, Tsao CY, Valdes JJ, Bentley WE. Cyclic AMP (cAMP) and cAMP receptor protein influence both synthesis and uptake of extracellular autoinducer 2 in *Escherichia coli*. J Bacteriol. 2005;187: 2066–2076. doi:10.1128/jb.187.6.2066-2076.2005

37. Pereira CS, Santos AJM, Bejerano-Sagie M, Correia PB, Marques JC, Xavier KB. Phosphoenolpyruvate phosphotransferase system regulates detection and processing of the quorum sensing signal autoinducer-2. Mol Microbiol. 2012;84: 93–104. doi:10.1111/j.1365-2958.2012.08010.x

38. Díaz-Pascual F, Lempp M, Nosho K, Jeckel H, Jo JK, Neuhaus K, et al. Spatial alanine metabolism determines local growth dynamics of *Escherichia coli* colonies. Elife. 2021;10. doi:10.7554/elife.70794

39. Velazquez EM, Nguyen H, Heasley KT, Saechao CH, Gil LM, Rogers AWL, et al. Endogenous Enterobacteriaceae underlie variation in susceptibility to *Salmonella* infection. Nat Microbiol 2019 46. 2019;4: 1057–1064. doi:10.1038/s41564-019-0407-8

40. Spragge F, Bakkeren E, Jahn MT, Araujo EBN, Pearson CF, Wang X, et al. Microbiome diversity protects against pathogens by nutrient blocking. Science. 2023;382(6676):eadj3502. doi:10.1126/science.adj3502

41. Leatham MP, Banerjee S, Autieri SM, Mercado-Lubo R, Conway T, Cohen PS. Precolonized human commensal *Escherichia coli* strains serve as a barrier to *E. coli* O157:H7 growth in the streptomycin-treated mouse intestine. Infect Immun. 2009;77: 2876–2886. doi:10.1128/iai.00059-09

42. Barthel M, Hapfelmeier S, Quintanilla-Martínez L, Kremer M, Rohde M, Hogardt M, et al. Pretreatment of mice with streptomycin provides a *Salmonella enterica* serovar Typhimurium colitis model that allows analysis of both pathogen and host. Infect Immun. 2003;71: 2839– 2858. doi:10.1128/iai.71.5.2839-2858.2003

43. Laganenka L, Colin R, Sourjik V. Chemotaxis towards autoinducer 2 mediates autoaggregation in *Escherichia coli*. Nat Commun. 2016;7: 12984. doi:10.1038/ncomms12984

44. Hegde M, Englert DL, Schrock S, Cohn WB, Vogt C, Wood TK, et al. Chemotaxis to the quorum-sensing signal AI-2 requires the Tsr chemoreceptor and the periplasmic LsrB AI-2-binding protein. J Bacteriol. 2011;193: 768–773. doi:10.1128/jb.01196-10

45. Stecher B, Barthel M, Schlumberger MC, Haberli L, Rabsch W, Kremer M, et al. Motility allows *S.* Typhimurium to benefit from the mucosal defence. CellMicrobiology. 2008. pp. 1166–1180. doi:10.1111/j.1462-5822.2008.01118.x

46. Taga ME, Miller ST, Bassler BL. Lsr-mediated transport and processing of AI-2 in *Salmonella* Typhimurium. Mol Microbiol. 2003;50: 1411–27. doi:10.1046/j.1365-2958.2003.03781.x

47. Miller ST, Xavier KB, Campagna SR, Taga ME, Semmelhack MF, Bassler BL, et al. *Salmonella* Typhimurium recognizes a chemically distinct form of the bacterial quorum-sensing signal AI-2. Mol Cell. 2004;15: 677–687. doi:10.1016/j.molcel.2004.07.020

48. Wotzka SY, Kreuzer M, Maier L, Arnoldini M, Nguyen BD, Brachmann AO, et al. *Escherichia coli* limits *Salmonella* Typhimurium infections after diet shifts and fat-mediated microbiota perturbation in mice. Nat Microbiol. 2019;4(12):2164–2174. doi:10.1038/s41564-019-0568-5

49. Hapfelmeier S, Ehrbar K, Stecher B, Barthel M, Kremer M, Hardt WD. Role of the *Salmonella* pathogenicity island 1 effector proteins SipA, SopB, SopE, and SopE2 in *Salmonella enterica* subspecies 1 serovar Typhimurium colitis in streptomycin-pretreated mice. Infect Immun. 2004;72: 795–809. doi:10.1128/iai.72.2.795-809.2004

50. Schubert C, Nguyen BD, Sichert A, Naepflin N, Sintsova A, Feer L, et al. Monosaccharides drive *Salmonella* gut colonization in a context-dependent manner. bioRxiv. 2024; 2024.08.06.606610. doi:10.1101/2024.08.06.606610

51. Xavier KB, Bassler BL. Regulation of uptake and processing of the quorum-sensing autoinducer AI-2 in *Escherichia coli*. J Bacteriol. 2005;187: 238–48. doi:10.1128/JB.187.1.238-248.2005

52. Maier L, Diard M, Sellin ME, Chouffane E-S, Trautwein-Weidner K. Granulocytes impose a tight bottleneck upon the gut luminal pathogen population during *Salmonella* Typhimurium colitis. PLoS Pathog. 2014;10: 1004557. doi:10.1371/journal.ppat.1004557

53. Nguyen BD, Cuenca V. M, Hartl J, Gül E, Bauer R, Meile S, et al. Import of aspartate and malate by DcuABC drives H2/Fumarate respiration to promote initial *Salmonella* gut-lumen colonization in mice. Cell Host Microbe. 2020;27: 922–936.e6. doi:10.1016/j.chom.2020.04.013

54. Yoo W, Shealy NG, Zieba JK, Torres TP, Baltagulov M, Thomas JD, et al. *Salmonella* Typhimurium expansion in the inflamed murine gut is dependent on aspartate derived from ROS-mediated microbiota lysis. Cell Host Microbe. 2024;32: 887–899.e6. doi:10.1016/j.chom.2024.05.001

55. Schubert C, Winter M, Ebert-Jung A, Kierszniowska S, Nagel-Wolfrum K, Schramm T, et al. C4-dicarboxylates and l-aspartate utilization by *Escherichia coli* K-12 in the mouse intestine: L-aspartate as a major substrate for fumarate respiration and as a nitrogen source. Environ Microbiol. 2021;23: 2564–2577. doi:10.1111/1462-2920.15478

56. Schubert C, Unden G. Fumarate, a central electron acceptor for Enterobacteriaceae beyond fumarate respiration and energy conservation. Adv Microb Physiol. 2023;82: 267–299. doi:10.1016/bs.ampbs.2022.10.002

57. Jones SA, Gibson T, Maltby RC, Chowdhury FZ, Stewart V, Cohen PS, et al. Anaerobic respiration of *Escherichia coli* in the mouse intestine. Infect Immun. 2011;79: 4218–4226. doi:10.1128/IAI.05395-11

58. Pereira FC, Berry D. Microbial nutrient niches in the gut. 2017;19: 1366–1378. doi:10.1111/1462-2920.13659

59. Freter R, Brickner H, Botney M, Cleven D, Aranki A. Mechanisms that control bacterial populations in continuous-flow culture models of mouse large intestinal flora. Infect Immun. 1983;39: 676. doi:10.1128/iai.39.2.676-685.1983

60. Freter R, Brickner H, Fekete J, Vickerman MM, Carey KE. Survival and implantation of *Escherichia coli* in the intestinal tract. Infect Immun. 1983;39: 686. doi:10.1128/iai.39.2.686-703.1983

61. Conway T, Cohen PS. Commensal and pathogenic *Escherichia coli* metabolism in the gut. Microbiol Spectr. 2015;3. doi:10.1128/microbiolspec.mbp-0006-2014

62. Conway T, Cohen PS. Applying the restaurant hypothesis to intestinal microbiota: Anaerobes in mixed biofilms degrade polysaccharides, sharing locally prepared sugars with facultative anaerobes that also colonize the intestine. Microbe. 2015;10: 324–328. doi:10.1128/microbe.10.324.1

63. Ross PA, Xu W, Jalomo-Khayrova E, Bange G, Gumerov VM, Bradley PH, et al. Framework for exploring the sensory repertoire of the human gut microbiota. MBio. 2024;15: e0103924. doi:10.1128/mbio.01039-24

64. Ortega Á, Zhulin IB, Krell T. Sensory repertoire of bacterial chemoreceptors. Microbiol Mol Biol Rev. 2017;81. doi:10.1128/mmbr.00033-17

65. Cherrak Y, Salazar MA, Yilmaz K, Kreuzer M, Hardt W-D. Commensal *E. coli* limits *Salmonella* gut invasion during inflammation by producing toxin-bound siderophores in a *tonB*-dependent manner. PLOS Biol. 2024;22(6): e3002616. doi:10.1371/journal.pbio.3002616

66. McCormick BA, Laux DC, Cohen PS. Neither motility nor chemotaxis plays a role in the ability of *Escherichia coli* F-18 to colonize the streptomycin-treated mouse large intestine. Infect Immun. 1990;58: 2957. doi:10.1128/iai.58.9.2957-2961.1990

67. Boulanger EF, Sabag-Daigle A, Thirugnanasambantham P, Gopalan V, Ahmer BMM. Sugar-phosphate toxicities. Microbiol Mol Biol Rev. 2021;85. doi:10.1128/mmbr.00123-21

68. Boulanger EF, Sabag-Daigle A, Baniasad M, Kokkinias K, Schwieters A, Wrighton KC, et al. Sugar-phosphate toxicities attenuate *Salmonella* fitness in the gut. J Bacteriol. 2022;204. doi:10.1128/jb.00344-22

69. Jeckelmann JM, Erni B. The mannose phosphotransferase system (Man-PTS) - Mannose transporter and receptor for bacteriocins and bacteriophages. Biochim Biophys Acta - Biomembr. 2020;1862: 183412. doi:10.1016/j.bbamem.2020.183412

70. Grandcí Ement C, Elanie Tannì Eres M, Moréra S, Moréra M, Dessaux Y, Faure D. Quorum quenching: role in nature and applied developments. FEMS Microbiol Rev. 2016;038: 86–116. doi:10.1093/femsre/fuv038

71. Dong YH, Wang LH, Zhang LH. Quorum-quenching microbial infections: mechanisms and implications. Philos Trans R Soc B Biol Sci. 2007;362: 1201. doi:10.1098/rstb.2007.2045

72. Datsenko KA, Wanner BL. One-step inactivation of chromosomal genes in *Escherichia coli* K-12 using PCR products. Proc Natl Acad Sci U S A. 2000;97: 6640–6645. doi:10.1073/pnas.120163297

73. Gaissmaier MS, Laganenka L, Herzog MKM, Bakkeren E, Hardt WD. The mobilizable plasmid P3 of *Salmonella enterica* serovar Typhimurium SL1344 depends on the P2 plasmid for conjugative transfer into a broad range of bacteria *in vitro* and *in vivo*. J Bacteriol. 2022;204. doi:10.1128/jb.00347-22/

74. Cherepanov PP, Wackernagel W. Gene disruption in *Escherichia coli*: TcR and KmR cassettes with the option of Flp-catalyzed excision of the antibiotic-resistance determinant. Gene. 1995;158: 9–14. doi:10.1016/0378-1119(95)00193-A

75. Porwollik S, Santiviago CA, Cheng P, Long F, Desai P. Defined single-gene and multi-gene deletion mutant collections in *Salmonella enterica* serovar Typhimurium. PLoS One. 2014;9: 99820. doi:10.1371/journal.pone.0099820

76. Daniel BBJ, Steiger Y, Sintsova A, Field CM, Nguyen BD, Schubert C, et al. Assessing microbiome population dynamics using wild-type isogenic standardized hybrid (WISH)-tags. Nat Microbiol. 2024;9: 1103–1116. doi:10.1038/S41564-024-01634-9

77. Datta S, Costantino N, Court DL. A set of recombineering plasmids for gram-negative bacteria. Gene. 2006;379: 109–115. doi:10.1016/j.gene.2006.04.018

78. Grant AJ, Restif O, Mckinley TJ, Sheppard M, Maskell DJ, Mastroeni P. Modelling within-host spatiotemporal dynamics of invasive bacterial disease. PLOS Biol. 2008;6(4):e74. doi:10.1371/journal.pbio.0060074

79. Sintsova A, Ruscheweyh H-J, Field CM, Feer L, Nguyen BD, Daniel B, et al. mBARq: a versatile and user-friendly framework for the analysis of DNA barcodes from transposon insertion libraries, knockout mutants, and isogenic strain populations. Bioinformatics. 2024;40(2):btae078. doi:10.1093/bioinformatics/btae078

