## Supplementary Figures S1-S4 for "Interplay between chemotaxis, quorum sensing, and metabolism regulates *Escherichia coli*-*Salmonella* Typhimurium interactions *in vivo*"

### Figures S1-S5

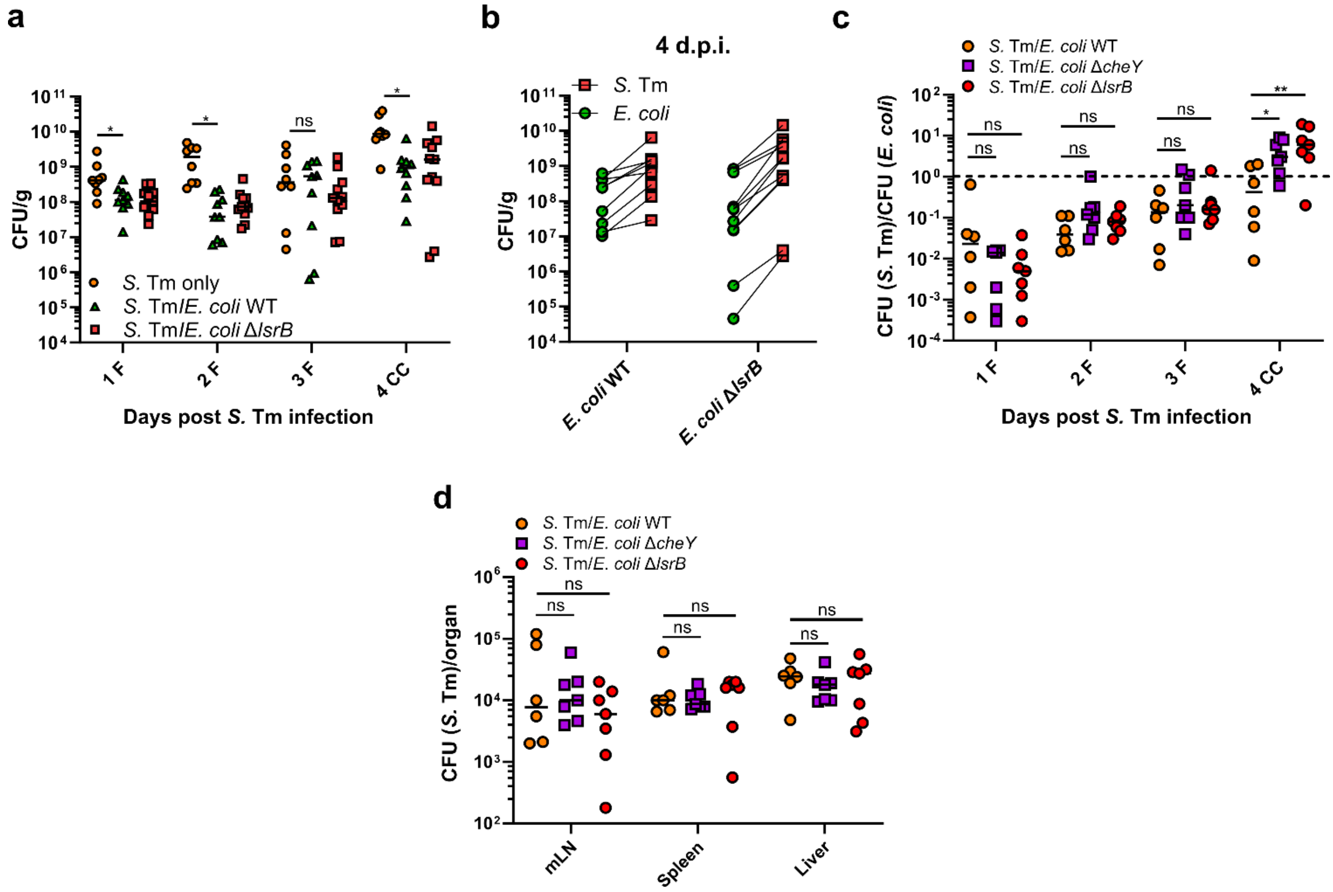


**Fig. S1. AI-2 chemotaxis-dependent *E. coli*-*S.* Tm competition in vivo. (a)** CFU counts of *S*. Tm in feces (F) and cecal content (CC) of mice infected with either *S.* Tm only or precolonized with *E. coli* wild type or Δ*lsrB* strains, as seen in Fig. 1b. The lines indicate median values (min mice n=8, at least two independent experiments). P values were calculated using the two-tailed Mann-Whitney *U*-test (*P<0.05, ns – not significant). **(b)** CFU counts of *E. coli* and *S.* Tm in mice precolonized with *E. coli* Z1331 wild-type or Δ*lsrB* at 4 days post *S.* Tm infection, as seen in Fig. 1b. **(c)** Competitive infections of *S.* Tm SL1344 against resident *E. coli* Z1331 wild-type, chemotaxis-deficient Δ*cheY* or AI-2 chemotaxis-negative Δ*lsrB* mutant strain in ampicillin-pretreated C57BL/6J SPF mice. The lines indicate median values (min mice n=6, at least two independent experiments). P values were calculated using the two-tailed Mann-Whitney *U*-test (**P<0.005, *P<0.05, ns – not significant). The dashed line indicates the CI value of 1. F, feces; CC, cecal content. **(d)** *S.* Tm counts in mesenteric lymph nodes (mLN), spleen and liver of *S.* Tm-infected mice as seen in panel (a). The lines indicate median values (min mice n=6, at least two independent experiments. P values were calculated using the two-tailed Mann-Whitney *U*-test (ns – not significant).

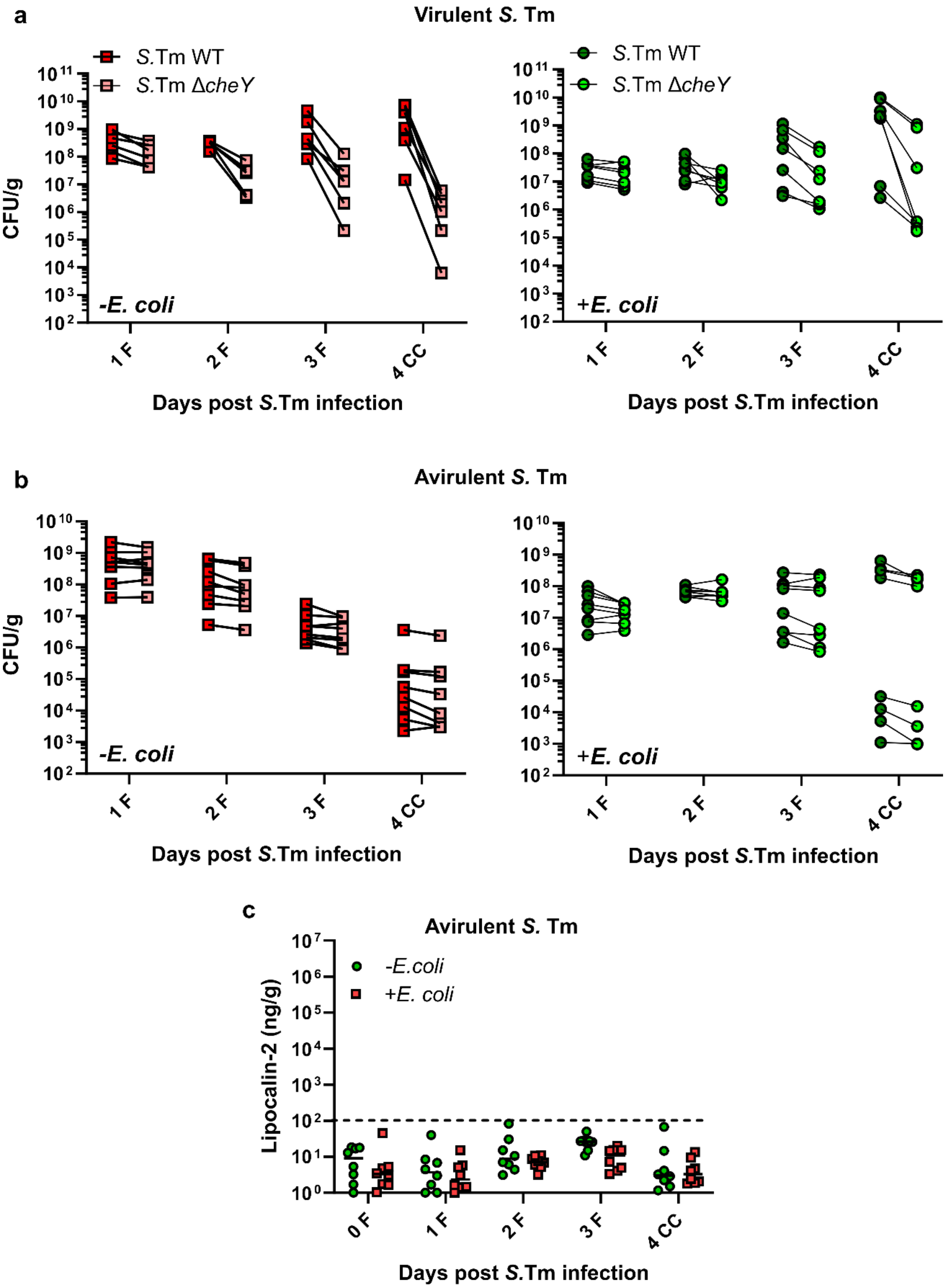


**Fig. S2. Inflammation-dependent fitness of *S.* Tm Δ*cheY*.** CFU counts of *S.* Tm SL1344 wild-type and chemotaxis-deficient Δ*cheY* strains in **(a)** virulent and **(b)** avirulent Δ*invG* Δ*sseD* background. Mice were either infected with *S.* Tm only or were precolonized with *E. coli* according to the experimental scheme shown in Fig. 1a. The gradual loss of CFU counts in avirulent *S.* Tm is due to its compromised ability to compete against the regrowing microbiota. **(c)** Lipocalin-2 levels in (F) feces and (CC) cecal content of mice infected with avirulent *S.* Tm SL1344 Δ*invG* Δ*sseD*. Dashed line indicates approximate level of lipocalin-2 marking a shift towards gut inflammation. Lines indicate median values (mice n=8, at least two independent experiments).

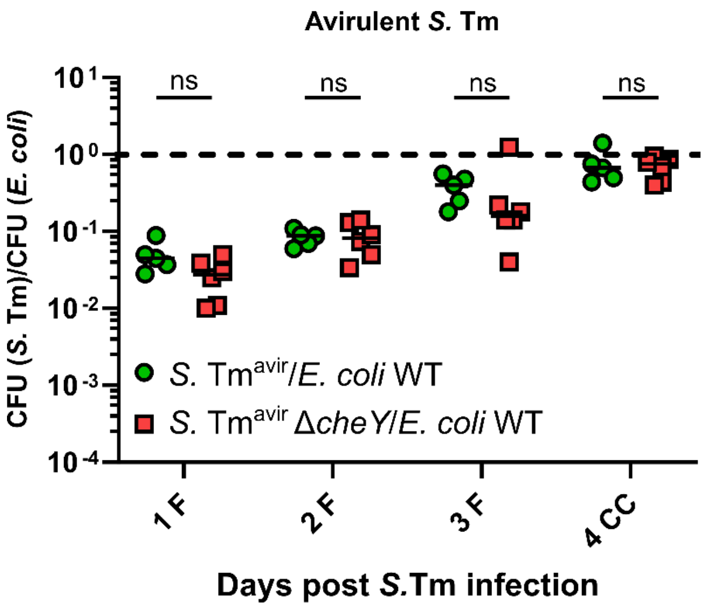


**Fig. S3. Chemotaxis is dispensable for *S.* Tm-*E. coli* competition in absence of *S*. Tm-induced inflammation.** Competitive infection of avirulent *S.* Tm SL1344 Δ*invG* Δ*sseD* strain (WT) and its non-chemotactic Δ*cheY* knockout strain against resident *E. coli* Z1331 strain. The lines indicate median values (min mice n=5, at least two independent experiments). P values were calculated using the two-tailed Mann-Whitney *U*-test (ns – not significant). The dashed line indicates the CI value of 1. F, feces; CC, cecal content.

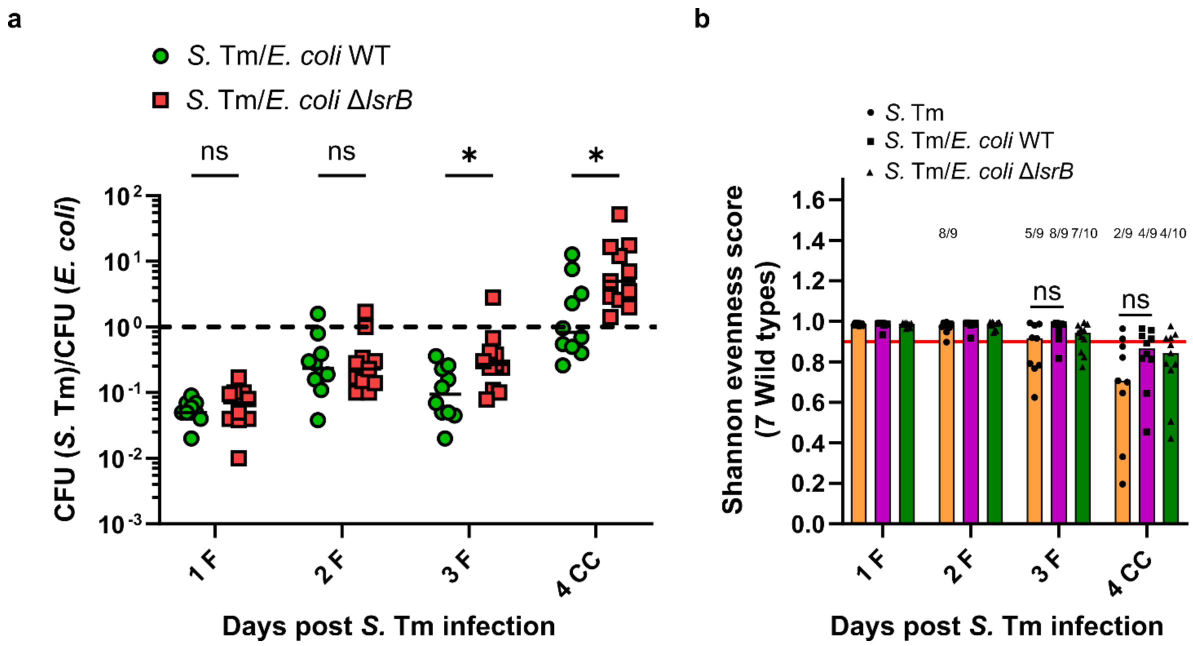


**Fig. S4. Validation of *S.* Tm SL1344 WISH-tagged wild-type and mutant pool. (a)** Competitive infections of *S.* Tm SL1344 WISH-tagged wild-type and mutant pool against resident *E. coli* Z1331 wild-type or AI-2 chemotaxis-negative Δ*lsrB* mutant strain. The lines indicate median values (min mice n=10, at least two independent experiments). P values were calculated using the two-tailed Mann-Whitney *U*-test (*P<0.05, ns – not significant). The dashed line indicates the CI value of 1. F, feces; CC, cecal content. **(b)** Shannon evenness score (SES) was calculated for the 7 WISH-barcoded SL1344 wild types. The red line indicates the SES of 0.9, which was the cutoff for further analysis. The number above the bar indicates how many samples are within this threshold.

| | *S.* Tm | | | | *S.* Tm + *E. coli* WT | | | | *S.* Tm +*E. coli* Δ*lsrB* | | | |
|---|---|---|---|---|---|---|---|---|---|---|---|---|
| | d1 | d2 | d3 | d4 | d1 | d2 | d3 | d4 | d1 | d2 | d3 | d4 |
| Δ*fruK* | 1.097 | 0.246 | 0.011 | 0.011 | 1.112 | 0.487 | 0.142 | 0.032 | 0.707 | 0.387 | 0.146 | 0.006 |
| Δ*srlD* | 0.905 | 0.836 | 0.683 | 0.447 | 0.564 | 0.742 | 0.613 | 0.724 | 0.853 | 1.169 | 1.113 | 1.667 |
| Δ*idnD* | 0.826 | 0.717 | 0.531 | 0.408 | 0.451 | 0.398 | 0.413 | 0.440 | 0.481 | 0.441 | 0.347 | 0.382 |
| Δ*glxK* | 1.013 | 1.079 | 1.436 | 1.890 | 1.060 | 1.046 | 1.046 | 0.860 | 1.231 | 1.108 | 0.982 | 0.791 |
| Δ*dgoK* | 0.067 | 0.046 | 0.019 | 0.026 | 0.448 | 0.430 | 0.394 | 0.715 | 0.582 | 0.422 | 0.274 | 0.374 |
| Δ*uxaC* | 0.800 | 0.888 | 0.206 | 0.084 | 0.798 | 0.824 | 0.891 | 0.887 | 1.136 | 1.280 | 1.151 | 0.340 |
| Δ*galK* | 0.546 | 0.335 | 0.256 | 0.376 | 0.844 | 0.698 | 0.760 | 0.613 | 0.871 | 0.831 | 0.796 | 0.562 |
| Δ*pgl* | 0.948 | 0.637 | 0.146 | 0.088 | 0.929 | 0.827 | 0.820 | 0.258 | 1.084 | 0.867 | 0.705 | 0.058 |
| Δ*rhaB* | 1.013 | 0.833 | 0.616 | 0.400 | 0.939 | 0.910 | 0.888 | 0.480 | 1.014 | 1.025 | 0.800 | 0.345 |
| Δ*malQ* | 0.881 | 2.022 | 6.931 | 7.385 | 0.803 | 1.046 | 1.553 | 3.421 | 0.899 | 1.239 | 1.332 | 1.965 |
| Δ*nagZ* | 1.039 | 1.234 | 1.166 | 1.094 | 0.884 | 1.003 | 0.907 | 1.117 | 1.057 | 1.110 | 1.395 | 0.810 |
| Δ*xylB* | 1.132 | 1.127 | 2.142 | 3.099 | 1.136 | 1.175 | 1.592 | 2.927 | 1.226 | 1.370 | 1.950 | 2.345 |
| Δ*manA* | 0.002 | 0.001 | 0.004 | 0.002 | 0.097 | 0.022 | 0.042 | 0.003 | 0.253 | 0.069 | 0.025 | 0.001 |
| Δ*gudD* | 0.999 | 1.218 | 4.508 | 6.747 | 0.687 | 0.702 | 0.792 | 1.668 | 0.788 | 0.885 | 1.088 | 2.678 |
| Δ*garK* | 1.252 | 1.174 | 0.943 | 0.858 | 1.289 | 1.427 | 1.697 | 2.092 | 1.587 | 1.482 | 1.591 | 0.893 |
| Δ*glpD* | 1.075 | 10.318 | 30.939 | 83.092 | 1.018 | 1.110 | 4.332 | 16.697 | 1.033 | 1.865 | 2.498 | 6.916 |
| Δ*araB* | 0.537 | 0.658 | 0.557 | 0.516 | 0.692 | 0.629 | 0.537 | 0.814 | 0.873 | 0.893 | 0.767 | 0.759 |
| Δ*nagB* | 1.089 | 0.925 | 0.333 | 0.765 | 0.795 | 0.774 | 0.249 | 0.270 | 0.825 | 0.502 | 0.115 | 0.010 |
| Δ*pfkB* | 1.390 | 1.881 | 2.568 | 4.078 | 1.102 | 1.370 | 1.591 | 3.449 | 1.378 | 1.253 | 1.358 | 2.238 |
| Δ*chbF* | 0.970 | 0.932 | 0.987 | 0.492 | 0.882 | 0.952 | 0.898 | 1.109 | 1.083 | 1.217 | 1.466 | 0.659 |
| Δ*gatD* | 0.948 | 0.935 | 0.796 | 0.674 | 0.782 | 0.928 | 0.922 | 0.903 | 1.031 | 0.976 | 0.924 | 0.624 |
| Δ*nagK* | 1.096 | 1.066 | 0.817 | 0.499 | 1.122 | 1.193 | 1.231 | 1.152 | 1.292 | 1.234 | 1.064 | 0.532 |
| Δ*pgm* | 0.002 | 0.000 | 0.001 | 0.001 | 0.003 | 0.001 | 0.001 | 0.002 | 0.002 | 0.001 | 0.002 | 0.000 |
| Δ*pgi* | 0.052 | 0.053 | LOD | LOD | 0.027 | LOD | LOD | LOD | 0.032 | 0.009 | 0.190 | LOD |
| Δ*mak* | 0.953 | 1.056 | 0.913 | 0.783 | 0.797 | 0.862 | 0.704 | 1.102 | 0.868 | 0.821 | 1.156 | 1.156 |
| Δ*nanK* | 0.951 | 0.918 | 0.703 | 0.384 | 0.817 | 0.818 | 0.910 | 1.175 | 1.137 | 0.898 | 1.201 | 0.720 |

fold change: 210, 5, 0.2, 0.001

| | *S.* Tm | | | | *S.* Tm + *E. coli* WT | | | | *S.* Tm +*E. coli* Δ*lsrB* | | | |
|---|---|---|---|---|---|---|---|---|---|---|---|---|
| | d1 | d2 | d3 | d4 | d1 | d2 | d3 | d4 | d1 | d2 | d3 | d4 |
| Δ*pfkA* | 0.049 | 0.022 | 0.001 | 0.028 | 0.015 | 0.009 | 0.003 | 0.003 | 0.025 | 0.009 | 0.006 | 0.002 |
| Δ*glpK* | 0.424 | 0.396 | 0.401 | 0.769 | 0.168 | 0.196 | 0.255 | 0.265 | 0.479 | 0.322 | 0.173 | 0.109 |
| Δ*treC* | 0.871 | 0.894 | 0.608 | 0.590 | 0.741 | 0.746 | 0.694 | 0.861 | 0.911 | 0.897 | 0.991 | 0.780 |
| Δ*fucK* | 1.083 | 1.146 | 0.858 | 0.988 | 1.062 | 1.003 | 1.057 | 1.006 | 1.258 | 1.062 | 1.070 | 0.854 |
| Δ*fbp* | 0.608 | 0.415 | 0.659 | 0.590 | 0.555 | 0.488 | 0.237 | 0.055 | 0.550 | 0.453 | 0.338 | 0.460 |
| Δ*tktA* | 0.021 | 0.001 | 0.001 | 0.024 | 0.041 | 0.014 | 0.005 | 0.034 | 0.125 | 0.034 | 0.017 | 0.000 |
| Δ*murQ* | 1.079 | 0.980 | 0.844 | 0.775 | 0.868 | 0.884 | 1.024 | 0.779 | 1.080 | 1.067 | 1.006 | 0.982 |
| Δ*edd* | 0.007 | 0.005 | 0.025 | 0.020 | 0.813 | 1.126 | 0.961 | 0.845 | 1.390 | 1.822 | 2.264 | 0.368 |
| Δ*tktB* | 2.459 | 1.659 | 2.547 | 53.887 | 3.558 | 3.371 | 5.345 | 70.062 | 2.952 | 2.284 | 6.207 | 1.248 |
| Δ*ppc* | 0.633 | 0.859 | 0.463 | 0.288 | 0.945 | 0.948 | 0.934 | 0.869 | 1.314 | 1.175 | 1.277 | 0.687 |
| Δ*kdgK* | 0.275 | 0.162 | 0.354 | 0.283 | 0.753 | 0.689 | 0.543 | 0.444 | 0.798 | 0.694 | 0.782 | 0.588 |
| Δ*ackA* | 0.005 | LOD | LOD | LOD | 0.035 | 0.010 | 0.005 | LOD | 0.051 | 0.016 | 0.039 | LOD |
| Δ*adhE* | 0.019 | 0.003 | 0.003 | 0.018 | 0.012 | 0.005 | 0.003 | 0.003 | 0.009 | 0.004 | 0.003 | 0.000 |
| Δ*gntK* | 0.434 | 0.328 | 0.129 | 0.097 | 0.694 | 0.590 | 0.432 | 0.444 | 0.743 | 0.737 | 0.521 | 0.243 |
| Δ*eda* | 0.133 | 0.020 | 0.002 | 0.005 | 0.125 | 0.072 | 0.019 | 0.005 | 0.218 | 0.038 | 0.006 | 0.001 |
| Δ*pflB* | 0.002 | 0.001 | 0.002 | 0.008 | 0.003 | 0.001 | 0.003 | 0.003 | 0.006 | 0.002 | 0.006 | 0.001 |
| Δ*glk* | 0.897 | 0.994 | 1.168 | 1.360 | 0.758 | 0.736 | 0.680 | 0.711 | 0.972 | 0.974 | 0.827 | 1.580 |
| Δ*glpA* | 0.965 | 0.974 | 0.480 | 0.438 | 0.776 | 0.644 | 0.531 | 0.267 | 0.974 | 0.633 | 0.511 | 0.433 |
| Δ*ptsG* | 1.160 | 1.090 | 0.857 | 0.334 | 1.227 | 1.183 | 1.108 | 0.908 | 1.529 | 1.209 | 1.182 | 0.860 |
| Δ*pckA* | 0.740 | 0.451 | 0.192 | 0.115 | 0.502 | 0.403 | 0.334 | 0.824 | 0.630 | 0.468 | 0.266 | 0.091 |
| Δ*ldhA* | 1.159 | 1.078 | 0.690 | 0.740 | 1.030 | 0.980 | 0.799 | 1.390 | 1.149 | 1.032 | 1.248 | 0.877 |
| Control-WT | 0.999 | 1.034 | 2.065 | 2.095 | 1.065 | 1.080 | 1.271 | 2.147 | 1.121 | 1.031 | 1.836 | 1.902 |
| Control-Δ*dcuABC* | 0.392 | 0.087 | 0.033 | 0.058 | 0.119 | 0.084 | 0.060 | 0.071 | 0.227 | 0.061 | 0.022 | 0.021 |
| Control-Δ*hyb* | 1.131 | 0.973 | 0.311 | 1.514 | 1.478 | 1.196 | 0.979 | 0.644 | 1.102 | 0.758 | 0.295 | 0.608 |
| Control-Δ*frd* | 0.292 | 0.050 | 0.011 | 0.018 | 0.082 | 0.031 | 0.019 | 0.105 | 0.129 | 0.025 | 0.012 | 0.006 |

**Fig. S5. Resident *E. coli* affects *S.* Tm central carbon metabolism during gut infection in both AI-2-dependent and -independent manners.** A heatmap showing the fitness of each *S.* Tm mutant in single infections and in competition with indicated *E. coli* strains. The shades of blue indicate loss of fitness, whereas the shades of red indicate gain of fitness, and white indicates a neutral effect. The CI values of all metabolic mutants tested are listed in Table S1. LOD, limit of detection as described in Materials and Methods.
